# No part gets left behind: Tiled nanopore sequencing of whole ASFV genomes stitched together using Lilo

**DOI:** 10.1101/2021.12.01.470769

**Authors:** Amanda Warr, Caitlin Newman, Nicky Craig, Ingrida Vendelė, Rizalee Pilare, Lilet Cariazo Cruz, Twinkle Galase Barangan, Reildrin G Morales, Tanja Opriessnig, Virginia Mauro Venturina, Milagros R Mananggit, Samantha Lycett, Clarissa YJ Domingo, Christine Tait-Burkard

**Affiliations:** The Roslin Institute and Royal (Dick) School of Veterinary Studies, University of Edinburgh, Easter Bush, Midlothian, UK; College of Veterinary Science and Medicine, Central Luzon State University, Science City of Muñoz, Nueva Ecija, The Philippines; Regional Animal Disease Diagnostic Laboratory, Department of Agriculture Regional Field Office III, City of San Fernando, Pampanga, The Philippines; Bureau of Animal Industry, Department of Agriculture, Visayas Avenue, Diliman, Quezon City, The Philippines

## Abstract

African Swine Fever virus (ASFV) is the causative agent of a deadly, panzootic disease, infecting wild and domesticated suid populations. Contained for a long time to the African continent, an outbreak of a particularly infectious variant in Georgia in 2007 initiated the spread of the virus around the globe, severely impacting pork production and local economies. The virus is highly contagious and has a mortality of up to 100% in domestic pigs. It is critical to track the spread of the virus, detect variants associated with pathology, and implement biosecurity measures in the most effective way to limit its spread. Due to its size and other limitations, the 170-190kbp large DNA virus has not been well sequenced with fewer than 200 genome sequences available in public repositories. Here we present an efficient, low-cost method of sequencing ASFV at scale. The method uses tiled PCR amplification of the virus to achieve greater coverage, multiplexability and accuracy on a portable sequencer than achievable using shotgun sequencing. We also present Lilo, a pipeline for assembling tiled amplicon data from viral or microbial genomes without relying on polishing against a reference, allowing for structural variation and hypervariable region assembly other methods fail on. The resulting ASFV genomes are near complete, lacking only parts of the highly repetitive 3’- and 5’telomeric regions, and have a high level of accuracy. Our results will allow sequencing of ASFV at optimal efficiency and high throughput to monitor and act on the spread of the virus.

## Main

African Swine Fever (ASF) is a viral hemorrhagic disease leading to an extremely high mortality of up to 100% within 7-10 days. Clinical signs of infection are non-specific, including fever, ataxia, anorexia, cyanosis, respiratory symptoms, gastrointestinal symptoms and death ^1^. The causative agent of ASF is African Swine Fever Virus (ASFV), the only member of the *Asfivirus* genus and the *Asfarviridae* family. The virion is large and complex with a diameter of around 175-215nm containing a large, double stranded DNA genome of around 170-190kb encoding over 150 open reading frames ^1,2^.

ASF is endemic in Africa where a sylvatic cycle between *Ornithodoros spp*. soft ticks and the natural reservoir, African wild suids, maintains its presence ^3^. Other routes of infection and spread include physical contact, fluids, excretions, contaminated feed and fomites. The virus is extremely resilient and can survive for prolonged periods in a range of environmental conditions in carcasses and pork product. The disease is highly contagious and can be transmitted through relatively low infectious dose in feed and water ^4^. Wild suid species are susceptible to disease but only domestic and feral pigs, as well as Eurasian wild boar show symptoms. ASFV is therefore very difficult to contain.

ASF was first discovered in East Africa, with symptoms reported in Kenya in 1914 and the disease described in 1921 ^5^. Outbreaks in other parts of Africa, Europe, Brazil and the Caribbean islands occurred in the 20^th^ century, with African countries being the worst affected. The virus was almost completely eradicated from non-African countries by the end of the century, but an outbreak in the Republic of Georgia in 2007 has since lead to widespread outbreaks in other countries. Since 2018 major outbreaks have been occurring in China, the world’s largest pork producer, and it has been reported that up to half of the pigs in the country, representing roughly a quarter of the world’s population, died or were culled to contain the outbreak in 2019 ^6^. The spread of the virus in East and Southeast Asia however could not be halted and since further countries including Mongolia, Vietnam, Cambodia, North Korea, Laos and island nations including The Philippines, Indonesia, Timor-Leste and Papua New Guinea have reported outbreaks ^7,8^. After the initial outbreaks in Eastern Europe and continuing spread, the disease reached the European Union in 2014 and continues to spread, not least through the wild boar population ^7,9^. In late 2020 the virus reached the largest producer of pork in Europe, Germany ^10^. The virus is also edging closer to the USA, one of the world’s main exporters and importers of pork, having been recently detected in Haiti and the Dominican Republic, only 381km by air from the US territory of Puerto Rico. It is clear, that the disease represents a serious panzootic threat impacting the pork industry and threatening economies already shaken by the SARS-CoV-2 pandemic.

To understand the genetic and genomic variation for ASFV, sequencing is primarily focused on a roughly 400bp fragment of the B646L gene, encoding for the major capsid protein p72. This fragment, representing <0.25% of the total genome, is the basis of the current genotyping system, which has identified 24 genotypes so far ^11–14^. To further discriminate, additional fragments of the E183L gene (p54, ~630bp), CP205L (p30, ~510bp), and B602L (gp83 ~800bp) are used, adding up to less than 1.4% of the genome characterized. Whilst being a DNA virus, antigenic diversity, the ability to acquire large deletions or insertions, and the presence of highly mutagenic hypervariable regions urge the need for whole genome sequencing for virus characterization and epidemiological studies ^15,16^. To do this at the scale required, there is a need for a cheap and efficient method to sequence the large ASFV genome, whilst high abundance of homopolymers and hypervariable region require highest accuracy ^17^.

The availability of a portable sequencing technology opens new doors to travel to outbreak locations, sequence, and analyze samples without needing to transport them. The MinION sequencer from Oxford Nanopore Technologies (ONT) can be carried easily in a pocket or carryon bag. This avoids complications of challenging transportation of biological samples, highly contagious agents or the requirement of a cold chain. There is the potential for a fast turnaround from sample collection to analysis, allowing for near live-monitoring of outbreak situations, as observed in during the Western African Ebola virus epidemic 2013-2016, or of course the COVID-19 pandemic. Furthermore, multiplexing and the washing and reuse of the most expensive component of sequencing, the flow cells, allowing for cheaper sequencing than other methods. Finally, the sequencer can produce very long reads which improves assembly potential, particularly of highly repetitive genomes.

Whilst it is possible to obtain whole genome sequences of ASFV directly from blood- and tissue extract DNA, the high prevalence of pig DNA and the need for baits or other methods to enrich ASFV DNA render that method inapplicable for high-throughput, fast, sequencing.

Here, we present a method to sequence the near complete genomes, excluding only the highly repetitive, variable length telomeric 3’ and 5’ regions, of ASFV using ONT’s MinION sequencing device using a tiled amplicon approach. The genome is amplified in 32 large fragments 7kb in length, amplified simultaneously in two PCR pools. We propose this method as an efficient, highly adaptable, more accurate, fast, and cost-effective option for sequencing of continuing ASFV outbreaks as well as historic samples. We present 10 complete ASFV genome assemblies from samples from the early stages of the ASFV outbreak in the Philippines in 2019 assembled either with the tiled sequencing approach or a whole genome sequencing shotgun approach. The portability of Nanopore sequencing makes it ideal for exploring the dynamics of ASFV infections as outbreaks emerge. As ASFV continues to spread around the world, efficient methods of sequencing the genome are essential to improve our understanding of the virus and the ongoing global spread. Our primer sets have been optimized for relatively even coverage and have been designed to bind outside of hypervariable regions. They only anneal to roughly 0.8% of the genome and are designed to be well suited to the current outbreak, able to at least partially sequence other genotypes and be easily modifiable should the virus mutate.

Finally, we present the Lilo pipeline. While pipelines exist to assemble genomes from tiled amplicons, they rely on aligning reads to a reference and using polishing tools to generate a consensus from the reads. This method works well for producing a genome sequence with SNPs representative of the sequenced genome, however large indels, structural variants, and hypervariable regions that may be difficult to align to a reference are not accurately represented. For ASFV, whole genes can be inserted or deleted and due to homologous recombination it can carry large structural variations, with indels likely being more important than SNPs in creating viral diversity ^18^. Therefore, we designed Lilo, which aligns reads to a reference in order to assign them to an amplicon, selects the read with the highest base quality and of the expected length for each amplicon, polishes the read with the remaining reads, removes primers and stitches them together at overlaps ordered and oriented by a reference. This approach makes the pipeline more adaptable to large structural variation and hypervariable regions in genomes than currently available methods.

## Shotgun sequencing of ASFV directly from blood

In field sequencing, particularly in developing countries, limits the availability of tools and reagents. During the first outbreaks in the Philippines whole DNA was isolated from the highly hemolysed blood collected from ASFV positive pigs. Samples were digested overnight with proteinase K at 55°C prior to phenol/chloroform/isoamyl alcohol extraction and precipitation with isopropanol before washing with 70% ethanol. Whole DNA samples were prepared for sequencing using the ligation sequencing kit (LSK) LSK109 before sequencing samples on a R9.4 flow cell using a MinION mk1b. The data were basecalled and demultiplexed using Guppy (ONT) and the reads assembled with Flye and polished with medaka. (Figure 1A)

**Figure 1.**
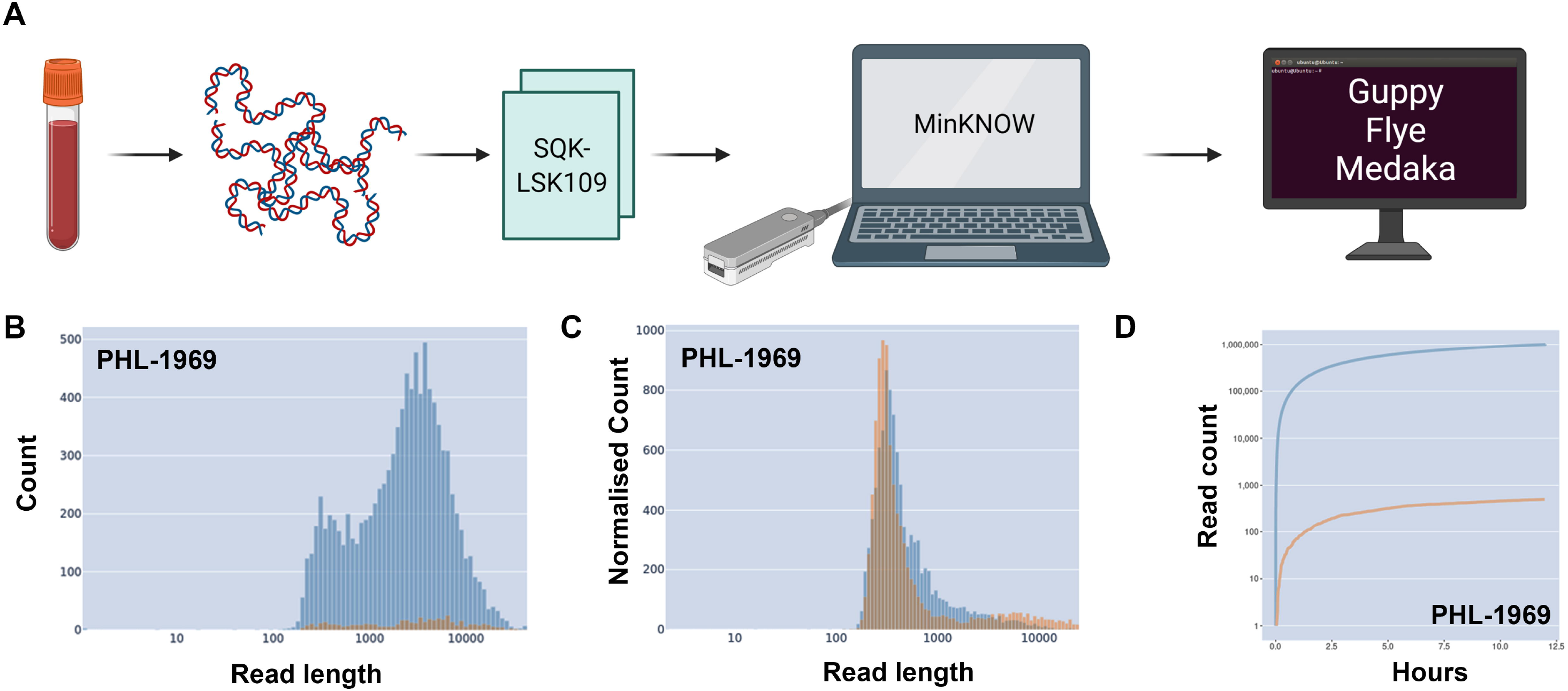
A) DNA was extracted from blood and sequenced with Nanopore’s LSK109 on an m1kb before analysis and assembly using Flye and polishing with medaka. B) Read length histograms for the dataset demonstrating total throughput (blue) and ASFV reads throughput (orange) C) Normalized counts by dataset of reads for total throughput (blue) and ASFV reads throughput (orange). D) Throughput over time for total read count (blue) and ASFV read count (orange).

The time between the beginning of sequencing and detection of the first ASFV read from whole blood ranged from 19 seconds to 3 minutes. As seen in the example of PHL-1969 (Figure 1B) the percentage of reads that came from ASFV ranged from 0.006% to 0.24%, likely dependent on the viral titers of the animals culled. ASFV samples show a similar size distribution to other DNA found in the samples, if anything a second small peak of larger fragments can be observed (Figure 1C). All four sequenced blood samples assembled into a whole genome, however, due to variable coverage, the number of mismatches and indels found in some of the samples were high (Figure 3B).

## Tiled amplicon sequencing of ASFV

Given the low yield of ASFV sequences from shotgun sequencing, as demonstrated by us and others ^19–21^, and the high expense per sample, this sequencing approach was not fit for purpose for high-throughput screening of an ongoing virus outbreak. Therefore, we developed a method to amplify, sequence, and assemble ASFV genomes from pigs.

In order to enrich ASFV from the sample easily, a PCR amplification approach was chosen, due to its ease of use and usually readily available tools in many countries and labs. Tiling primers were designed targeting 7kb amplicon length and 1kb amplicon overlap using primal scheme using a set of 26 ASFV reference sequences (Figure 2A). The primers are well suited to genotype II, from the current outbreak, but also cover the majority of the genome for at least genotypes I and IV (Figure 2B). This relatively long amplicon size was chosen to reduce the number of primer pairs but also to span potential hypervariable regions. After initial individual performance tests, several primers were redesigned from the original set of primers produced by primal scheme, however the majority of them worked well from the beginning. Fragments were amplified using the PCRBio VeriFi Hot Start high fidelity polymerase according to the manufacture’s instruction. Following redesign, all primers amplified their targets, however, they did so at different efficiencies leading to uneven coverage over the genome. To test this, evenly concentrated pools of primers (pool 1 and pool 2, Figure 2A and Figure 2C) were used to amplify blood DNA extract samples from ASFV-infected pigs. Following initial amplifications, pools were split into three pools with primer pair 1, producing a shorter 4kb fragment continuously outperforming the others in a mixed reaction on its own, and primer concentrations in pool 1(Pair 1) and pool 2 were gradually adjusted according to their performance. PCR products per sample were combined, libraries prepared using the LSK109 kit in an R9.4 flow cell. Figure 2D demonstrates the improvement that can be gained by tweaking primer concentrations from evenly represented primer pairs (purple) to optimized primer concentrations (green). These optimizations improve performance for multiplexing of multiple samples on one flow cell.

**Figure 2.**
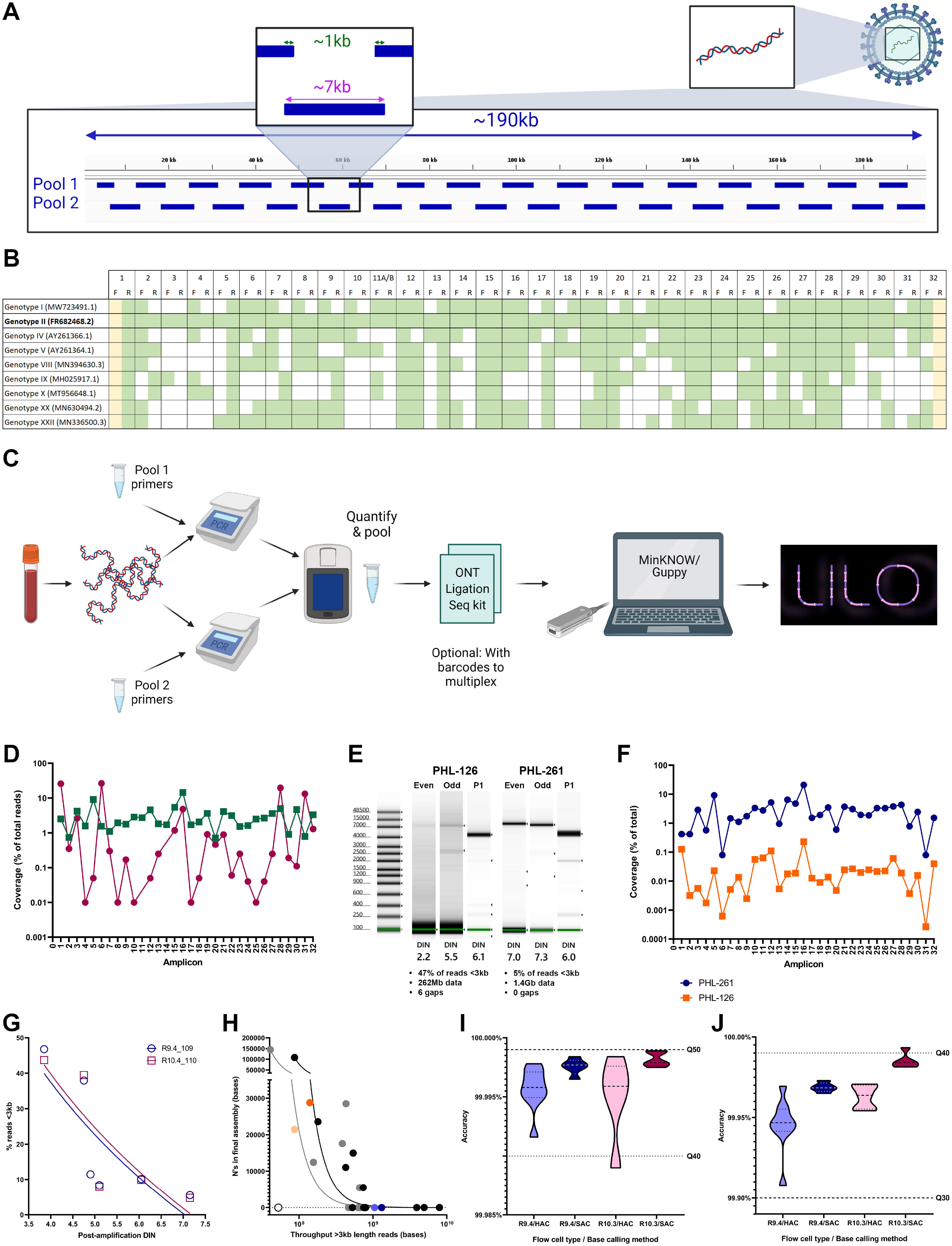
A) Design of the tiled primer scheme for ASFV with ~7kb amplicons and ~1kb overlaps. B) Predicted primer binding in correct region for representative ASFV genotypes. C) Workflow with extraction from blood, PCR amplification of primer pools, pooling, sequencing and bioinformatic analysis D) Coverage of one sample (PHL-3142) amplified with either evenly represented primer pairs (purple) or optimized proportions of primer pairs (green). E) Tapestation capillary electrophoresis of amplified pools of two different samples, the low quality PHL-126 and the high quality PHL-261, and statistics of the resulting assemblies of each of these. F) Coverage of amplicons using optimized primer concentrations for the two samples from figure 1E. G) Impact of post-amplification DIN on proportion of reads <3kb in length, essentially wasted sequencing capacity, using R9.4 flow cells and LSK-109 (magenta) or R10.4 and LSK-110 (blue). Semilog fit analysis shows an R squared correlation of 0.6420 or 0.7244, for R9.4_109 and R10.4_110, respectively. H) Association between overall throughput >3kb and number of gaps in final assembly using HAC (black/bold) and SAC (grey/faint). A log-log analysis shows an R squared correlation of 0.9680 and 0.9358 for HAC and SAC, respectively. PHL-126 is highlighted in orange and PHL-261 in blue. I) Assembly accuracy based on proportion of mismatches against reference (MN715134.1), with lines showing Q40 and Q50 PHRED scores. J) Assembly accuracy based on proportion of indels against reference (MN715134.1), with lines showing Q30 and Q40 PHRED scores. A&C Created with BioRender.com

**Figure 3.**
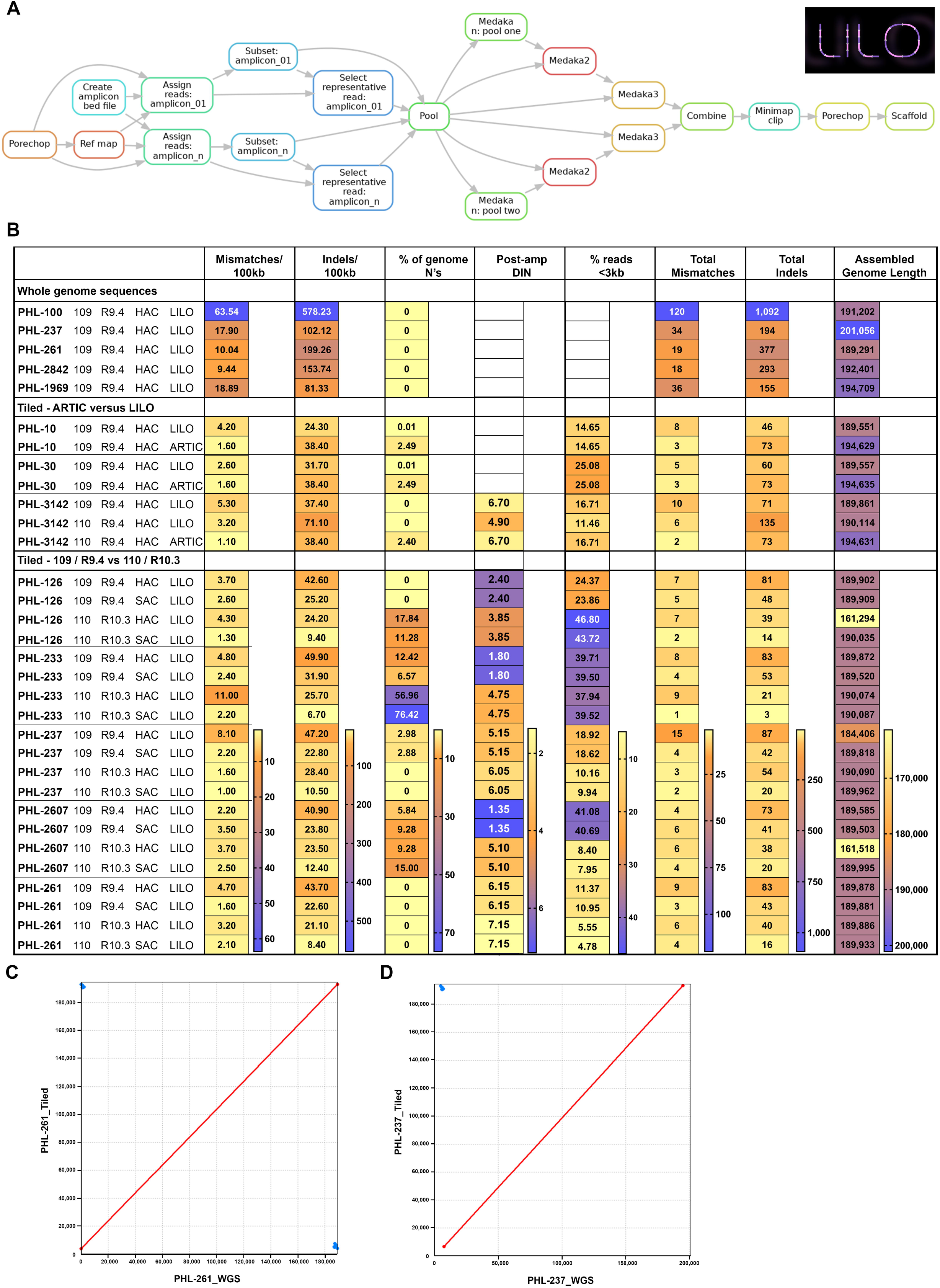
A) Directed acyclic graph showing the steps the Lilo pipeline takes during assembly. The graph has been simplified to show assembly of a genome containing 2 amplicons (amplicon_01 and amplicon_n) for a single sample. B) Quast results for genomes sequenced with tiled amplicons or from shotgun sequencing on R9.4 flow cells using SQK-LSK109. Note PHL-10 and PHL-30 were sequenced with an earlier version of the primer scheme with one primer different and are expected to have a 25bp gap C) Nucmer alignment of PHL-261 genomes assembled from WGS or Lilo tiled assembly. D) Nucmer alignment of PHL-237 genomes assembled from WGS or Lilo tiled assembly.

Fresher samples amplify more cleanly, but older, degraded samples will still amplify sufficiently. To show this, we highlight two samples; sample PHL-126, which has been heavily used and degraded, and sample PHL-261, which has been used less frequently (aliquot stored in freezer without frequent use) and is of better quality. As can be seen in the automated electrophoresis result of a tapestation (Figure 2E), PHL-126 shows poor amplification and relatively many amplicons <7kb. Good amplification can be seen for the shorter amplicon pair 1 still. PHL-261 on the other hand shows continued good amplification of the desired 7kb and 4kb products of pool 1 (odd), pool 2 (even) and pair 1, respectively. These samples were prepared with the LSK109 kit and multiplexed using native barcoding and run on a R9.4 flow cell with 3 other ASFV genomes having been pooled in representative quantities, the poorer amplification of PHL-126 had lower sequencing throughput than the better quality PHL-261, but was still assembled into a near-complete genome. Figure 2F shows sequencing coverage of the same two samples and the proportion of total reads for each that was assigned to each of the 32 amplicons.

The post-amplification DNA Integrity Number (DIN) can be used to help predict multiplexability, as different quality samples will impact the needed throughput. Figure 2G shows the relationship between the post-amplification DIN and throughput <3kb, and Figure 2H demonstrates the number of gaps for different throughputs of reads >3kb. Samples PHL-126 (orange) and PHL-261 (blue) have been highlighted in pale (super accuracy base calling (SAC)) and strong (high accuracy base calling (HAC)) colors, respectively. Figure 2H demonstrates the required throughput per sample and the gaps that can be expected in genomes produced with lower throughputs. Assuming a theoretical MinION throughput of 30GB, it should be possible to multiplex over 24 samples, and potentially up to 48 samples (including PCR control), however we would recommend starting with fewer and assessing achievable throughput for each sequencing location as there is variability in expected throughput between users, flow cells, and geographical location. The integrity of the sample will also impact the throughput with degraded samples leading to sequencing capacity being taken up by shorter fragments instead of the required full length amplicons (Figure 2G). For very poor samples, more stringent size selection with AMPure XP beads prior to sequencing may be necessary if samples are to be multiplexed. While it is impractical to run an automated, high resolution electrophoresis, such as a tapestation, after every amplification, users can test a typical sample type (e.g. from decomposing wild boar, from farm culled pigs, with/without cold chain) to predict the likely multiplexability and clean-up steps of similar samples.

## Lilo assembly of genomes from tiled amplicons

Comparing ASFV genomes we found major variation of the genome often originating from indels. Available assembly pipelines were struggling with such variation when it did not correspond to the reference sequence. Therefore, we developed the Lilo pipeline (Figure 3A) to assemble the tiled amplicons (https://github.com/amandawarr/Lilo). Whilst Lilo uses a reference alignment to sort the amplicons, it polishes against the highest quality reads rather than a reference sequence. Using this pipeline, highly accurate genomes were obtained with mismatch accuracy approaching Q50 when using SAC (Figure 2I) and indel accuracy up to Q40 when compared to a closely related publicly available ASFV genome assembly (MN715134.1) ^21^, which may still be quite divergent from these samples in truth.

QUAST ^22^ (v5.0.2; quality assessment tool for genome assemblies) results demonstrate that the increased coverage of the tiled amplicons produced a more accurate assembly than shotgun sequencing of the virus using a whole flow cell sequencing directly from extracted DNA. Shotgun sequencing however, was able to highlight some samples with longer telomeric regions, such as PHL-237, which is a clear advantage of long-read sequencing technology and something that should be explored for more in-detail investigations into the role of the ASFV telomeric regions. Overall, SAC produced fewer mismatches and indels than HAC and should be the preferred method, however, the time for base calling is a trade-off. Samples with high percentages of unassigned bases (N’s) clearly correspond to DIN numbers (Figure 3B).

The assembled genomes had excellent agreement on genome structure with the same samples assembled from shotgun sequencing (Figures 3B & 3C). The repetitive content shown at the edges of Figures 3B and 3C are sequences from the telomeres, showing that despite the sequences being from tiled amplicons, they do cover the majority of the genome and part of the telomeres.

## Accuracy of Lilo and ARTIC assembled genomes

We assessed the quality of Lilo assemblies against those produced with the ARTIC pipeline (v1.2.1). A selection of the ASFV sequencing data were assembled using the ARTIC pipeline, as well as using Lilo, both using the assembled shotgun sequence PHL-1969 as a reference.

QUAST analysis shows lower numbers of mismatches against the closest reference (MN715134.1) but higher indels. The percentage of unassigned bases is much higher for ARTIC at around 2.4% whereas Lilo is at 0 or nearly 0%.(Figure 3B)

Comparing Lilo-assembled genomes and ARTIC-assembled genomes to a reference (MN715134.1) a number of indels can be observed. Figure 4A shows a likely real indel in the PHL ASFV samples which all assemblies agree on and which is well supported by the reads. In contrast, Figure 4B shows the only indel unique to almost all of the assemblies produced by the Lilo pipeline while being absent from all artic assemblies and occurs in a homopolymer, Most reads appear to support the deletion assembled by Lilo, whether this is a real sequence or a result of poor accuracy of Nanopore sequencing of homopolymeric regions is a more difficult question. Figure 4C shows an extreme example of a very long homopolymeric region, ASFV has several of these and typically neither assembly method agrees on the length of the homopolymer, with the reads lending no strong support to either assembly. While errors from the Lilo pipeline tended to be randomly dispersed among homopolymers, ARTIC errors tended to be more systematic, appearing consistently across the assembled genomes. Frequently, homopolymers lead to the ARTIC pipeline replacing the base immediately before the homopolymer and the first base of the homopolymer with a pair of N’s, as can be seen in Figure 4E and 4F. There were also occasions when reads did not support an indel, and there wasn’t a clear cause of an indel in the reads or reference, as can be seen in Figures 4D and 4E. There are several of these types of indels throughout the assembly, where Lilo assemblies better agree with both the reads and the reference, likely contributing to the lower QUAST scores of ARTIC on homopolymers and the percentage of undefined bases.

**Figure 4.**
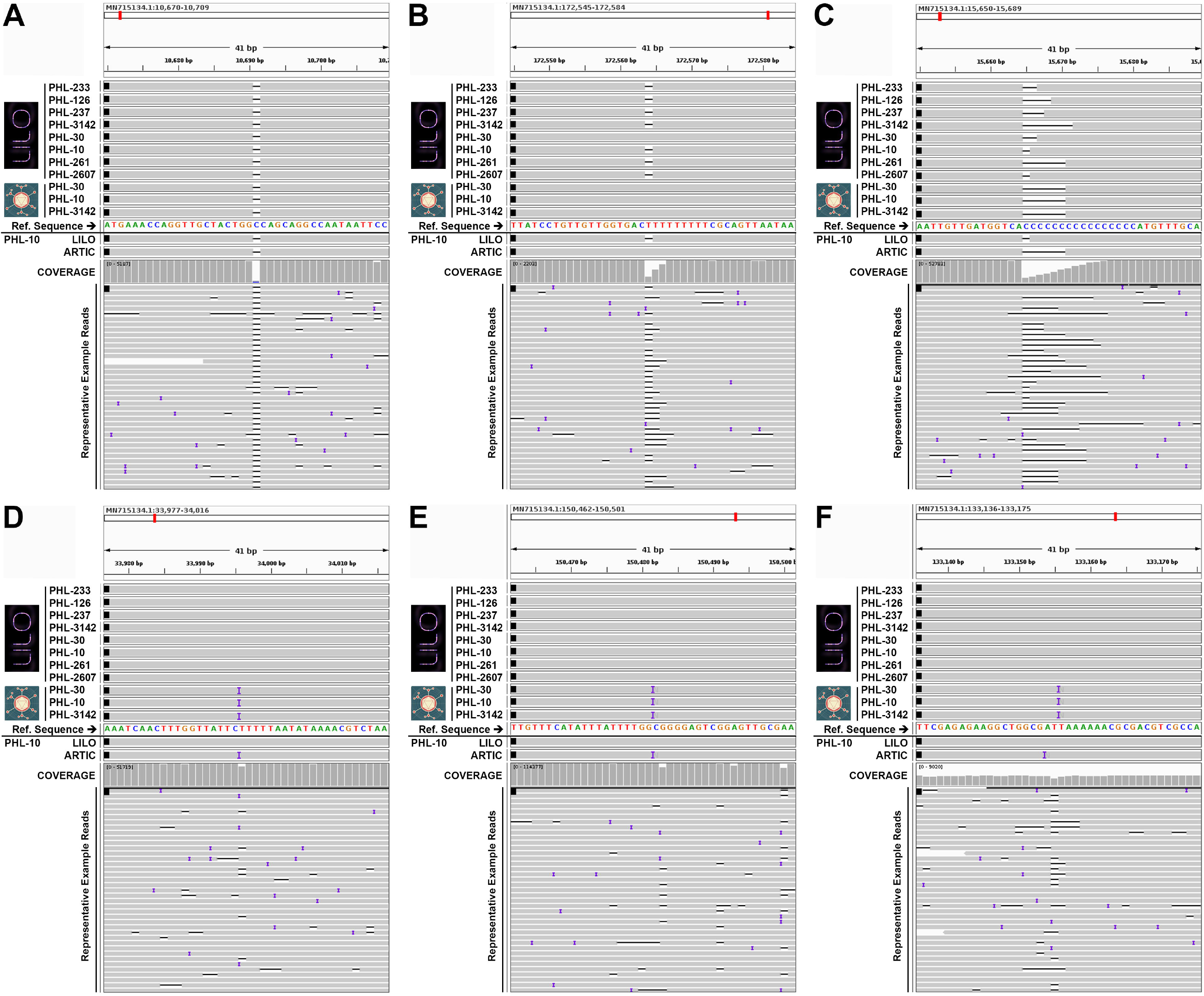
B) IGV image showing alignment of genomes assembled with Lilo or Artic (top) and assemblies and reads for a single sample (PHL-10) aligned to a reference (MN715134.1). This image shows a likely real indel present in all assemblies and supported by the reads. C) as in B, but showing an indel common in Lilo assemblies and missing in Artic assemblies and the reference. D) as in B, but showing a long homopolymer with poor consensus from the reads and inconsistent results in assemblies. E, F & G) Examples of indels specific to the Artic pipeline assemblies which do not agree with the reference.

## Phylogenomics

A maximum likelihood phylogenomic tree was constructed incorporating the newly assembled genomes from tiled data of R9.4 flow cells with HAC with publicly available whole genome ASFV sequences using iqtree (v2.0.5; Figure 5A), and for B646L gene, encoding for the major capsid protein p72, genotype II sequences specifically (Figure 5B).

**Figure 5.**
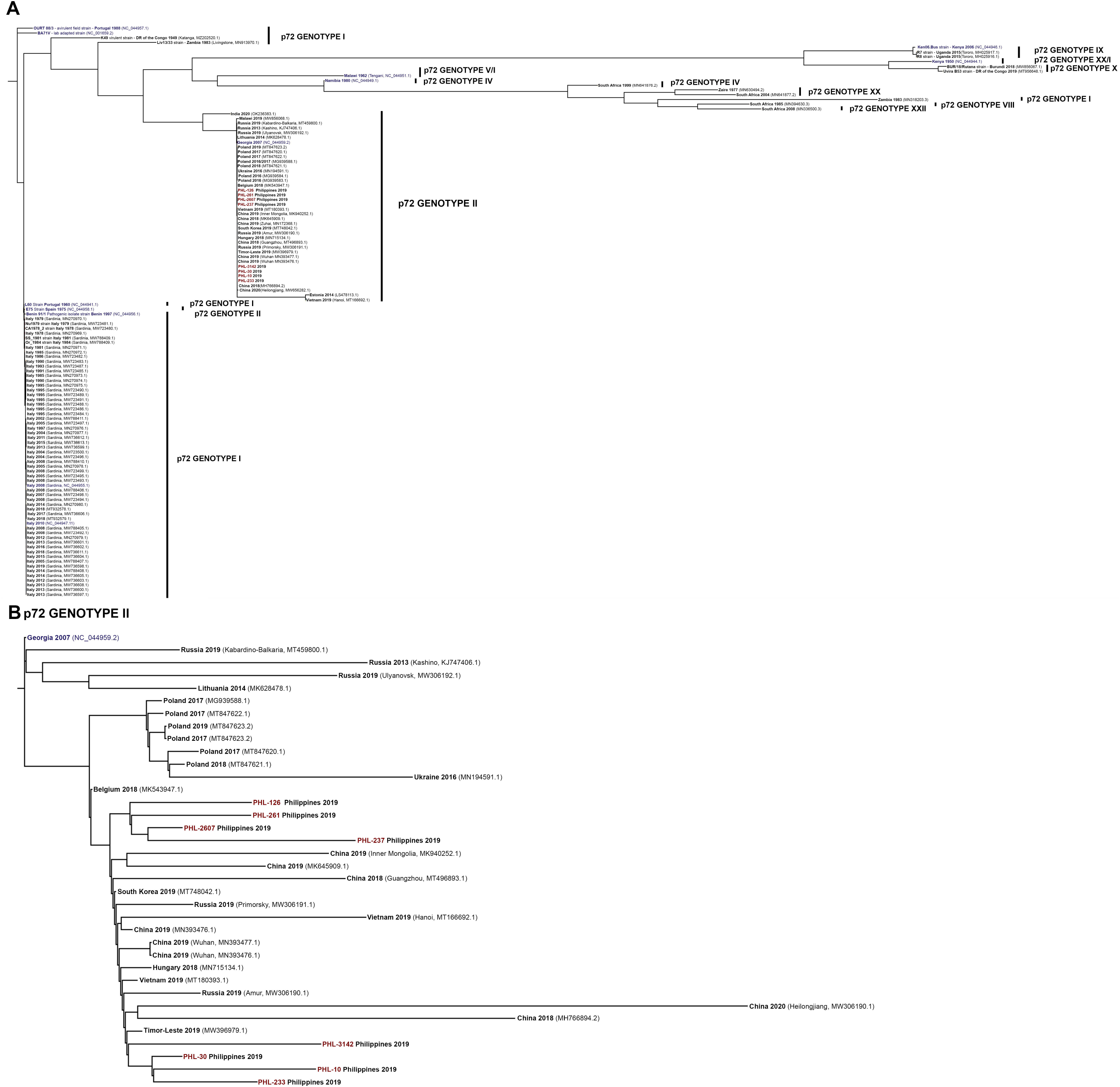
Maximum likelihood trees for our R9.4/SQK-LSK109 genomes and A) all available ASFV genomes downloaded from NCBI (09/11/2021) or B) those specifically clustering with genotype II.

As observed in Figure 5A, p72 genotypes do not correspond to the clustering. For example, the E75 strain Spain 1975 isolate, an early genotype II, is grouping with genotype I’s. Unfortunately, the phylogeny contains many gaps and lacks both timely and geographic resolution, showing that much more sampling is required. PHL samples clearly cluster within the highly virulent, novel p72 genotype II cluster.

Resolving the tree further, selecting only those clustering with the novel p72 genotype II genomes two distinct clusters of PHL sequences can be observed. Whilst, due to the similarity of the genomes, the orders of lower branches are of lower confidence than those of higher branches, reanalysis still suggest two different introductions into the Philippines (internal branch lengths may be found in supplementary documents S1 and S2). Whilst indications are that one cluster is closer related to Asian isolates, the other one showing a more likely Eastern European origin, the lack of sample numbers to fill branch gaps and common ancestors makes conclusive interpretation impossible.

## Discussion

ASFV is a serious threat to the global pork industry and consequences of depopulation, reduced availability of pork, and increased prices of other animal protein affect local economies, especially in low- and middle income countries with high reliance on pig protein. Tracing the spread of the virus, understanding more about genome-pathology links, and consequently implementing targeted biosafety measures are paramount to combat the disease. As demonstrated in Figure 5, current genotyping methods based on partial sequencing of the B646L gene, encoding for the major capsid protein p72, do not correspond to whole-genome sequencing and are therefore inadequate to trace virus evolution.

Here, we have demonstrated an efficient method for sequencing the ASFV genome. Despite the use of tiled amplicons, part of the 3’- and 5’ telomeric regions of the virus are included in the tiled amplicon assemblies, meaning the majority of the genome is included.

As demonstrated by us and others, ASFV sequences can be obtained by direct sequencing from blood or other tissue samples of infected pigs ^19–21^. The resulting sequence includes interesting information on the lengths and repeats found in the telomeric regions, which may be helpful for more in-depth investigation into the virus pathology and spread. However, without enrichment for ASFV ^17^ or depletion of host-methylated DNA ^23^ sample percentage for ASFV is low relative to host DNA in the samples, meaning that obtaining sufficient ASFV reads to assemble the genome from shotgun sequencing usually requires an entire MinION flow cell, or more, depending on viral titer and original sample type. Bone marrow or blood will likely yield the best virus:host ratio with spleen or muscle, whilst good sources of viral DNA ^24^, also contain a large number of nucleated host cells. Even if sufficient data is obtained to assemble the genome, the coverage is likely too poor to sufficiently polish the genome. In contrast, the tiled amplicon method can be used on samples with lower viral titers or degraded DNA, selectively sequences the virus, and can be multiplexed on a flow cell to simultaneously sequence multiple samples at high enough coverage for good polishing. Especially in countries where ASFV is circulating in wild boar or feral pigs, samples may be collected from infected animals that have been dead for a prolonged period of time. It is important that the method is capable of amplifying virus from both high- and low quality samples. Figure 2E demonstrates the variability of DNA integrity post-amplification and that even poor samples that have been degraded amplify and produce near complete genome assemblies.

Overall, the PCR amplification method increases coverage, is less prone to exhaust flow cells quickly, allows for multiplexing, and consequently reduces costs, improves genome accuracy, and removes the need for specialized enrichment or depletion methods.

Whilst ~7kb amplicons are very large compared to other comparable methods for other viruses, the size of the ASFV genome, the stability of DNA, the relatively low numbers of primer pairs, and the advantages of long reads detecting recombinants more easily make this the best approach. Especially with the small, medium, and large indels that can occur in ASFV ^18^, it is important to get good resolution across these regions, which can be achieved easily with large amplicons. It is important though to choose the right, high accuracy polymerase capable of amplifying such long amplicons. We found PCRBio VeriFi to be highly capable of this with the hot start version producing very few non-specific products, whilst the non-hot start version can produce more non-specific product, which may be an advantage for variant testing. As demonstrated in Figures 2E and F and 4B show that even low quality samples can produce whole genome assemblies with few gaps. However, a limitation of the large tiled amplicon method is that should a variant occur at the site of a primer, the amplification of a relatively large section of genome will fail. While this is an inconvenience, it will be simple to redesign a primer to replace the failed one or to act as an alternate primer. It is also possible to amplify across a larger region using the existing primers either side of the failed one, generating a 14kb product, to sequence a larger region and design a primer from the sequenced amplicon. This was found to be possible using the VeriFi HS polymerase and allows for the method to adapt as the virus changes.

Whilst Nanopore sequencing methods provide a lot of advantages, such as sequencing on site, portability, and accessibility to less specialist communities, there are, as for any sequencing method, drawbacks.

As demonstrated in Figure 1I & 1J the choice of flow cell and basecalling method has an impact on the accuracy of SNPs and indels. Generally, throughput is lower and required input DNA is higher for R10.3 flow cells, however HAC on R10.3 is of comparable accuracy to SAC on R9.4, with super accuracy on R10.3 being of even higher quality. SAC is very slow and resource hungry, and when speed is important such as in an active outbreak, R10.3 with HAC may be a good compromise for maximising efficiency with minimal sacrifice of accuracy. However where the highest accuracy is required we recommend using R10.3 flow cells and basecalling with a super accuracy model.

All of our assemblies have indels compared to the reference partially stemming from systematic errors in Nanopore sequencing in the abundant homopolymers and repeats in the ASFV genome (e.g. PHL-1969 contains 6.8% homopolymers of 4bp or more), however with the latest kits and flow cells from Nanopore, and without additional costs, we would expect future developments to continually reduce the indel errors without major alterations to the wet or dry lab methodologies described here. Additionally, some indels are likely true variations, and in many cases these indels are clearly present across all of our genomes (Figure 4A), and are not in homopolymeric regions, suggesting true variation between samples and the reference used. Additional sequencing of the same amplicons with a higher accuracy technology, such as Illumina could be used to polish the assemblies, however this would add time and cost. Reducing the number of false indels, where possible, is important, as the ASFV genome is known to have functionally relevant indels ^25^. Polishing with an accurate reference can produce assemblies that are very accurate, however, these methods do not handle structural variants and hypervariable regions well. While the genomes sequenced here do not have any major indels compared to the reference used, diversity in ASFV is partially driven by small, medium and large indels ^18^ and increased sequencing of samples is likely to reveal more of them. While errors from the Lilo pipeline tended to be randomly dispersed among homopolymers, ARTIC errors tended to be more systematic, appearing consistently across the assembled genomes. Errors occurring in the same position between genomes may be more likely to impact phylogenomic analysis than relatively random errors. The only consistent indel error found across the majority of the Lilo assembled genomes that was always absent in the artic genomes is shown in Figure 4B. This region contains a homopolymers, which is typically difficult to correct from Nanopore sequencing data, however while the ARTIC assembly more closely agrees with the reference, the reads are well-supporting of the deletion found in the Lilo assemblies. It is not unusual when carrying out multi-sequence alignments between whole ASFV genome sequences, even those constructed from reads from a higher accuracy sequencing technology, to find large homopolymers of variable length and it is unclear to what degree these are limitations of sequencing technologies as opposed to real variation.

The Lilo pipeline also has some limitations, it currently assumes that any structural variants will not change the length of any given amplicon by more than 5%, it assumes that structural variants will not be dramatic enough to prevent alignment to the reference for the purposes of assigning reads to amplicons and ordering and orienting the polished amplicons. Lilo also assumes the reads will be the full length of the amplicon, making it incompatible with ONT rapid kits that utilize transposases. However, the strength of not relying on polishing reads aligned to a reference is beneficial for genomes where structural variation is expected to be important, and for species with hypervariable regions which may not align and polish well with a reference. The pipeline has been tested on tiled sequences from ASFV, Porcine Reproductive and Respiratory Syndrome-1 & −2, and SARS-CoV-2 (data not shown here) and can handle custom schemes for other viruses.

ASFV is very under-sequenced with only a small number of whole genome sequences available and there is a need for an affordable way to sequence the virus at scale, as previously discussed ^17^. While the majority of sequences produced from sequencing individual genes in the virus have been frustratingly similar ^26^ reducing their usefulness in epidemiological studies, variants and large deletions have been observed across the genome, and these have been found to affect phenotypes ^25^. As outbreaks continue to spread around the world and the amount of virus in circulation increases, these variations will likely increase in frequency. Additionally, there are few of the ancestral viruses from Africa sequenced, and these should be sequenced to understand the evolution of ASFV, particularly the loss of its dependence on the sylvatic cycle. Given the slow mutational rate of the virus, sequencing individual genes is unlikely to be informative and so to have a chance of seeing variants in the virus the whole genome must be sequenced. The ability to amplify the genotypes with our current scheme decreases with distance from genotype II, and additional primers will need designing in the future to improve coverage over other genotypes, however current coverage using this primer scheme is still likely to be of more use than the p72 gene alone. Coverage gaps can be resolved relatively easily as larger amplicons can be generated with flanking primers. Should primers on older or emerging samples fail, the altered region can be amplified using primers from either side of the failed amplicon, spanning the region, and the sequenced amplicon can be used to design new primers for the region.

Phylogenetic trees of currently available whole-genome ASFV sequences highlight the inadequacy of the p72 genotyping in reflecting similarities of ASFV on a genomic level. The sparsity of whole-genome sequences hampers the ability to trace virus movement and results in high levels of uncertainty in phylogenetic analyses. Running the maximum likelihood tree analysis including the whole-genome sequences obtained from tiled amplification in this manuscript reliably grouped samples from the early Philippines outbreak of ASFV in November 2019 into two clusters. This indicates at least two potential introductions of ASFV into the Philippines. Due to the lack of samples and resolution in the phylogenetic tree, no conclusion about countries or region of origins is possible.

We have presented an efficient, low cost method for sequencing and assembling ASFV which can be carried out in the lab or in the field during outbreaks. The Lilo pipeline is a lightweight pipeline that can be run on a standard laptop with 16GB RAM and no internet connection, making it ideal for in field bioinformatic analysis of ASFV and other viruses.

## Methods

### Samples

Blood samples from outbreaks in central Luzon (Philippines) were collected following depopulation of pigs within a defined containment radius. Blood samples were tested for ASFV by PCR. Blood samples from ASFV-positive pigs were pooled at equal amounts by farm before further processing.

### DNA extraction

Blood samples were spun for 20min at 3,000rcf before decanting the supernatant. 5xTEN buffer(0.05M EDTA, 0.5M NaCl, 20mg/ml Proteinase K, 20% SDS, in 0.05M Trix-HCl, pH8.0) were added to a 1x final concentration before incubation overnight at 55°C in a shaking water bath. Equal volumes of phenol were added and gently mixed. Following 20min centrifugation at 3,000rcf the aqueous phase was transferred to a fresh tube. If the phase was very viscous, the phenol phase was re-extracted to improve yields. An equal volume of phenol/chloroform/isoamyl alcohol (25:24:1) was added to the aqueous phase before mixing and separation by centrifugation, 10min, 3,000rcf. The aqueous phase was transferred to a fresh tube before addition of 1:10 3M sodium acetate and an equal amount of isopropanol. Following 1h incubation at −20°C, samples were spun for 10min at 16,000rcf before washing the pellet with 70% Ethanol. The pellet was dried and resuspended in nuclease-free water.

### Nanopore sequencing directly from DNA extracted from blood in the Philippines

Samples were sequenced following Nanopore’s SQK-LSK109 protocol on R9.4 flow cells on a MinION mk1b. The protocol was started with 1ug of DNA as measured on an Implen NanoPhotometer P330. The protocol was carried out as recommended by ONT with the following modifications: The 20°C and 60°C incubations after the addition of NEB’s FFPE repair and End-prep reagents were done for 30 minutes at each temperature instead of 5 minutes, and the room temperature incubation for the ligation reaction was done for 20 minutes instead of 10. During sequencing, two USB desk fans were pointed at the MinION to assist with maintaining appropriate temperature for the run in the above average “room temperature” in the lab in the Philippines.

### Designing primers for tiled amplification

Tiled primers were designed using Primal Scheme (v1.3.2) ^27^. A set of 28 complete African Swine Fever genomes (listed in Supplementary Document S3) were downloaded from NCBI for primer design, which at the time were all that were available. Additionally three Filipino whole ASFV genomes we had assembled from shotgun sequencing data were included. A multi sequence alignment was carried out with Clustal Omega (in MEGA v7.0.2) ^28^. Primal Scheme was run to produce 7kb amplicons with 1kb overlap resulting in 32 overlapping primer pairs in two non-overlapping pools. Primers were tested on samples and while the majority worked first time, several had to be redesigned due to failed amplification or preferential amplification of off-target regions. Redesigns were done using Primer-BLAST ^29^, targeting a similar region of the genome to the failed amplicon. Some primers amplified more efficiently than others and in order to make the coverage of these as even as possible, some primers were tweaked to have a different concentration. One primer pair (amplicon 1) was shorter than the others in order to avoid highly repetitive sequence in the telomeres and it is recommended to amplify it in a separate reaction to pools 1 and 2 to avoid overrepresentation.

### Amplification, library prep and sequencing of tiled amplicons

Tiled primers were initially tested individually at 200nM concentration using approximately 90ng ASF DNA and Phusion High-Fidelity PCR Master Mix with HF Buffer with 1mM added MgCl_2_ (both New England Biolabs, Ipswich, MA, USA). Individual PCRs, in a 25μl reaction volume, underwent initial denaturation of 2 minutes at 98°C, followed by 33 cycles of 10 seconds at 98°C, 30 seconds annealing at 63°C, and 4 minutes and 40 seconds extension at 72°C, followed by a final extension for 10 minutes at 72°C. PCR products were then examined using Tapestation Genomic DNA analysis (Agilent, Santa Clara, CA, USA). While a small amount of off-target amplification was tolerated, primers which produced strong off-target bands or weak bands of the correct 7kb size were redesigned.

Once the complete set of primers had been successfully designed to cover the complete genome, the primers were pooled in equal amounts into two pools of non-overlapping primers. These pools were tested using the same conditions as the individual PCRs, but in a 50μl reaction volume and using 1μM of the primer pool. The resulting PCR products were cleaned using 0.4× volume AMPure XP beads (Beckman Coulter, Indianapolis, IN, USA) to remove products smaller than approximately 2kb in length, then pooled equally prior to sequencing. The cleaned PCR products were quantified using a Qubit ds DNA BR assay (Invitrogen, Waltham, MA, USA) and combined in equimolar amounts to a total of 700ng for library preparation according to the Native barcoding genomic DNA (with EXP-NBD104, EXP-NBD114, and SQK-LSK109)-Nanopore protocol.

Following bioinformatic analysis of sequencing data, primers which were found to be over- or under-performing were either redesigned or their contribution to the pool was adjusted accordingly, and the new primer pool tested as above in an iterative fashion. Ultimately 2 non-overlapping pools and a separate reaction for primer pair 1 were used to obtain the most even coverage and were processed as above, and pooled proportionally to the number of amplicons in each pool prior to sequencing. Additionally the polymerase was swapped from Phusion to VeriFi (PCRBIO) in a 25ul reaction using 2ul DNA per reaction, which has markedly better performance on the amplicons with far less off-target amplification. The PCR conditions for this polymerase were an initial denaturation of 1 minute at 98°C, followed by 40 cycles of 15 seconds at 98°C, 15 seconds annealing at 60°C, and 4 minutes and 40 seconds extension at 72°C, followed by a final extension for 5 minutes at 72°C. AMPure XP bead cleanup after PCR is optional, but recommended in samples with low DIN. Primer sequences, recommended primer concentrations and recommended pooling quantities are described in supplementary table S1, and any updates to these will be released on Lilo’s github page.

Samples were sequenced following Nanopore’s SQK-LSK109 or SQK-LSK110 protocol on R9.4 or R10.3 flow cells (which combination is specified alongside relevant results) on a MinION mk1b or mk1c. The protocol was started with 1ug of pooled amplicons as measured on a qubit using broad range reagents. For samples using multiplexing, the native barcoding expansion kit from Nanopore was used following Nanopore’s instructions when using SQK-LSK109. For using the barcodes with SQK-LSK110, the instructions for SQK-109 were followed until after the barcodes had been ligated on, at which stage the end prep was repeated and we follow the standard protocol for library prep with SQK-LSK110 from after the end prep step.

### Bioinformatic processing of ASFV genomes sequenced with shotgun sequencing

The data were basecalled and demultiplexed using MinKNOW (v19.06.8; ONT) using “fast” basecalling. Following basecalling the reads were aligned to an ASFV genome using minimap2 to identify ASFV reads, the fast5s for these reads were extracted using fast5_subset from the ont_fast5_api (https://github.com/nanoporetech/ont_fast5_api) and these were basecalled again using high accuracy basecalling. This was done to reduce basecalling time, as this work was done locally in the field on a laptop without a GPU. The reads were assembled with Flye (v2.6) ^30^ and polished 3 times with Medaka (v0.7.1; ONT). Comparisons of quantity of data produced and the proportion of which were ASFV reads were done using NanoComp (v1.28.1) ^31^.

### Bioinformatic processing of ASFV genomes from tiled amplicons with Lilo

The data were basecalled and demultiplexed using Guppy (v5.0.14; ONT) using high or super accuracy model on a GPU. The snakemake pipeline, Lilo (https://github.com/amandawarr/Lilo), was developed and as summarised in Figure 3A, takes the following steps:

1. Use Porechop (v0.2.3) to remove any sequencing adapters or barcodes that have made it through demultiplexing.
2. Align to a reference with minimap2 (v2.22) ^32^ and samtools (v1.12) ^33^ and separate reads into amplicons by alignment position with bedtools (v2.30.0).^34^
3. Select reads of the expected amplicon length (+/-5%) and subset to 300X
4. Select the read with highest average base quality within +/-1% of the median length of reads for the amplicon to be the “reference” (with bioawk v1); https://github.com/lh3/bioawk), remove any amplicons with fewer than 40 reads. Targeting the median length allows for flexibility for large insertions or deletions.
5. Pool amplicon reads and references back into their original non-overlapping pools.
6. Polish the pools 3x with medaka (v1.4.4; ONT) and combine resulting polished amplicons.
7. Align to the reference with minimap2 and remove soft clipped bases (these likely represent missed barcodes or adapters)
8. Run porechop (specific fork: https://github.com/sclamons/Porechop-1) to remove primers from the amplicons.
9. Merge the amplicons with scaffold_builder (v2.3) ^35^.

The required input to Lilo are demultiplexed reads in fastq format in a directory named “raw/”, a reference fasta, a bed file of primer alignments (as output by primal scheme), and a csv of primer sequences (if there are ambiguous bases it is advised to expand them first) and a config file, described on the github page. It is adaptable to any species (with a single genome fragment/chromosome) with any tiled primer scheme. The pipeline outputs a fasta file containing the assembled genome.

### ARTIC assemblies

A subset of genomes were also assembled using the Artic pipeline (https://artic.network/ncov-2019; v1.2.1) following the bioinformatics SOP using the medaka method.

### Quality control of assembled genomes

Quast (v5.0.2) was used to compare the assembled genomes to the most closely related publicly available ASFV assembly according to BLAST alignment (MN715134.1) ^21^. Samples where both WGS and tiled sequencing were used were compared for overall structure using nucmer (v4.0.0beta2) ^36^.

### Phylogeny

The phylogeny analysis was limited to the tiled genomes, as these were the most accurate assemblies, and publicly available genomes. These were aligned using Mafft (v7.467)^37^ and maximum likelihood trees constructed using iqtree (v2.0.5)^38^.

## Acknowledgments

We would like to thank the Bureau of Animal Industry and Milagros Mananggit for providing us access to the valuable ASFV blood samples. We would also like to thank Central Luzon State University and Clarissa Yvonne Domingo and Virginia Venturina and their families for their amazing hospitality during all our visits to the Philippines.

We acknowledge financial support from the BBSRC Institute Strategic Programme grant funding to The Roslin Institute (BBS/E/D/20241866, BBS/E/D/20002172, and BBS/E/D/20002174) and BBSRC / Newton Fund Swine and Poultry research initiative grant (BB/R013187/1).

## Author Contributions

Outlined the study AW, NC, IV, CYJD, CTB; collected and provided samples RGM, MRM, CYJD; performed experiments AW, NC, CN, IV, RP, LCC, TGB, and CTB; analyzed data AW, NC, SL, and CTB; interpreted data AW, NC, CN, SL, VMV, CYJD and CTB; wrote manuscript, AW and CTB, with contribution of other authors.

## Competing Interests Statement

The authors declare no competing interests.

## Notes

### Competing Interest Statement

The authors have declared no competing interest.

