## Supplementary for "No part gets left behind: Tiled nanopore sequencing of whole ASFV genomes stitched together using Lilo": Supplementary Document S1_tree details for 5A.docx

Supplementary document X- Output from iqtree2 detailing model selection and full likelihood statistics for the tree in figure 5A.

IQ-TREE 2.0.5 built May 15 2020

Type of analysis: ModelFinder + tree reconstruction

Random seed number: 31758

REFERENCES

----------

To cite IQ-TREE please use:

Bui Quang Minh, Heiko A. Schmidt, Olga Chernomor, Dominik Schrempf,

Michael D. Woodhams, Arndt von Haeseler, and Robert Lanfear (2020)

IQ-TREE 2: New models and efficient methods for phylogenetic inference

in the genomic era. Mol. Biol. Evol., in press.

https://doi.org/10.1093/molbev/msaa015

To cite ModelFinder please use:

Subha Kalyaanamoorthy, Bui Quang Minh, Thomas KF Wong, Arndt von Haeseler,

and Lars S Jermiin (2017) ModelFinder: Fast model selection for

accurate phylogenetic estimates. Nature Methods, 14:587–589.

https://doi.org/10.1038/nmeth.4285

SEQUENCE ALIGNMENT

------------------

Input data: 123 sequences with 208630 nucleotide sites

Number of constant sites: 168787 (= 80.9026% of all sites)

Number of invariant (constant or ambiguous constant) sites: 168787 (= 80.9026% of all sites)

Number of parsimony informative sites: 31606

Number of distinct site patterns: 20773

ModelFinder

-----------

Best-fit model according to BIC: GTR+F+R2

List of models sorted by BIC scores:

Model	LogL	AIC	w-AIC	AICc	w-AICc	BIC	w-BIC

GTR+F+R2	-644917.829 1290329.658 - 0.000913 1290330.246 - 0.000917 1292860.993 +	0.963

GTR+F+R3	-644908.831 1290315.663 +	0.999 1290316.260 +	0.999 1292867.494 -	0.0373

TVM+F+R2	-644940.719 1290373.438 - 2.84e-13 1290374.021 - 2.86e-13 1292894.524 - 5.04e-08

TVM+F+R3	-644932.060 1290360.120 - 2.22e-10 1290360.713 - 2.22e-10 1292901.703 - 1.39e-09

TIM+F+R2	-645035.755 1290561.510 - 4.12e-54 1290562.088 - 4.16e-54 1293072.348 - 1.23e-46

TIM+F+R3	-645027.553 1290549.106 - 2.03e-51 1290549.694 - 2.04e-51 1293080.440 - 2.14e-48

K3Pu+F+R2	-645056.750 1290601.500 - 8.53e-63 1290602.074 - 8.63e-63 1293102.089 - 4.27e-53

K3Pu+F+R3	-645048.550 1290589.100 - 4.2e-60 1290589.684 - 4.23e-60 1293110.187 - 7.44e-55

GTR+F+I+G4	-645398.129 1291290.259 - 2.34e-212 1291290.847 - 2.35e-212 1293821.593 - 2.46e-209

TIM2+F+R2	-645467.262 1291424.523 - 1.64e-241 1291425.102 - 1.65e-241 1293935.361 - 4.87e-234

TIM2+F+R3	-645459.083 1291412.167 - 7.88e-239 1291412.755 - 7.92e-239 1293943.501 - 8.31e-236

TPM2+F+R2	-645491.143 1291470.286 - 1.89e-251 1291470.860 - 1.91e-251 1293970.875 - 9.45e-242

TPM2u+F+R2	-645491.143 1291470.286 - 1.89e-251 1291470.860 - 1.91e-251 1293970.875 - 9.45e-242

TPM2+F+R3	-645483.173 1291458.346 - 7.4e-249 1291458.929 - 7.45e-249 1293979.432 - 1.31e-243

TPM2u+F+R3	-645483.191 1291458.381 - 7.27e-249 1291458.965 - 7.32e-249 1293979.468 - 1.29e-243

TN+F+R2	-645643.286 1291774.571 - 1.59e-317 1291775.145 - 1.61e-317 1294275.161 - 7.96e-308

TN+F+R3	-645635.016 1291762.031 - 8.41e-315 1291762.614 - 8.47e-315 1294283.117 - 1.49e-309

TIM3+F+R2	-645643.468 1291776.936 - 4.88e-318 1291777.514 - 4.92e-318 1294287.774 - 1.45e-310

TIM3+F+R3	-645634.993 1291763.986 - 3.16e-315 1291764.574 - 3.18e-315 1294295.320 - 3.34e-312

HKY+F+R2	-645664.395 1291814.790 -	0 1291815.359 -	0 1294305.131 - 2.47e-314

HKY+F+R3	-645656.173 1291802.347 - 1.48e-323 1291802.925 - 1.48e-323 1294313.185 - 4.41e-316

TPM3u+F+R2	-645664.334 1291816.667 -	0 1291817.241 -	0 1294317.257 - 5.75e-317

TPM3+F+R2	-645664.334 1291816.667 -	0 1291817.241 -	0 1294317.257 - 5.75e-317

TPM3u+F+R3	-645656.136 1291804.272 - 4.94e-324 1291804.855 - 4.94e-324 1294325.358 -	1e-318

TPM3+F+R3	-645656.142 1291804.283 - 4.94e-324 1291804.866 - 4.94e-324 1294325.369 - 9.96e-319

GTR+F+G4	-646333.107 1293158.214 -	0 1293158.797 -	0 1295679.300 -	0

GTR+F+I	-647019.719 1294531.438 -	0 1294532.021 -	0 1297052.524 -	0

SYM+R2	-651777.869 1304043.738 -	0 1304044.312 -	0 1306544.328 -	0

SYM+R3	-651770.457 1304032.913 -	0 1304033.496 -	0 1306553.999 -	0

TVMe+R2	-651811.322 1304108.644 -	0 1304109.213 -	0 1306598.986 -	0

TVMe+R3	-651803.468 1304096.935 -	0 1304097.513 -	0 1306607.773 -	0

TIMe+R2	-652626.732 1305737.463 -	0 1305738.028 -	0 1308217.556 -	0

TIMe+R3	-652619.316 1305726.631 -	0 1305727.205 -	0 1308227.221 -	0

TIM2e+R2	-652636.671 1305757.343 -	0 1305757.907 -	0 1308237.436 -	0

TIM2e+R3	-652629.190 1305746.379 -	0 1305746.953 -	0 1308246.969 -	0

K3P+R2	-652661.876 1305805.752 -	0 1305806.312 -	0 1308275.597 -	0

K3P+R3	-652654.457 1305794.914 -	0 1305795.483 -	0 1308285.255 -	0

TIM3e+R2	-653199.285 1306882.570 -	0 1306883.135 -	0 1309362.663 -	0

TIM3e+R3	-653191.854 1306871.707 -	0 1306872.281 -	0 1309372.297 -	0

TNe+R2	-653493.973 1307469.947 -	0 1307470.506 -	0 1309939.791 -	0

TNe+R3	-653486.882 1307459.763 -	0 1307460.332 -	0 1309950.104 -	0

K2P+R2	-653529.857 1307539.713 -	0 1307540.269 -	0 1309999.310 -	0

K2P+R3	-653522.770 1307529.541 -	0 1307530.105 -	0 1310009.633 -	0

F81+F+R2	-657501.658 1315487.317 -	0 1315487.881 -	0 1317967.409 -	0

F81+F+R3	-657494.883 1315477.767 -	0 1315478.340 -	0 1317978.356 -	0

JC+R2	-664576.369 1329630.739 -	0 1329631.289 -	0 1332080.087 -	0

JC+R3	-664569.996 1329621.993 -	0 1329622.553 -	0 1332091.837 -	0

GTR+F	-664817.233 1330124.465 -	0 1330125.044 -	0 1332635.303 -	0

AIC, w-AIC	: Akaike information criterion scores and weights.

AICc, w-AICc : Corrected AIC scores and weights.

BIC, w-BIC	: Bayesian information criterion scores and weights.

Plus signs denote the 95% confidence sets.

Minus signs denote significant exclusion.

SUBSTITUTION PROCESS

--------------------

Model of substitution: GTR+F+R2

Rate parameter R:

A-C: 1.4704

A-G: 6.0430

A-T: 2.1592

C-G: 2.0139

C-T: 6.5899

G-T: 1.0000

State frequencies: (empirical counts from alignment)

pi(A) = 0.3053

pi(C) = 0.1936

pi(G) = 0.1923

pi(T) = 0.3088

Rate matrix Q:

A	-0.9005	0.1213	0.4951	0.2841

C	0.1912	-1.223	0.165	0.8671

G	0.7859	0.1661	-1.084	0.1316

T	0.2808	0.5436	0.08193	-0.9063

Model of rate heterogeneity: FreeRate with 2 categories

Site proportion and rates: (0.886,0.3278) (0.114,6.222)

Category Relative_rate Proportion

1	0.3278	0.886

2	6.222	0.114

MULTIPLE RUNS

-------------

Run	logL

1	-644825.1806

2	-644890.4834

3	-644829.6411

4	-644827.8280

5	-644892.4990

6	-644827.8400

7	-644830.9600

8	-644826.8872

9	-644829.9612

10	-644829.1779

11	-644828.7023

12	-644830.8965

13	-644829.0533

14	-644890.5820

15	-644828.8383

16	-644828.4622

17	-644828.5438

18	-644828.2439

19	-644828.0255

20	-644890.5230

MAXIMUM LIKELIHOOD TREE

-----------------------

Log-likelihood of the tree: -644825.1806 (s.e. 25236314721.8293)

Unconstrained log-likelihood (without tree): -1061990.5783

Number of free parameters (#branches + #model parameters): 247

Akaike information criterion (AIC) score: 1290144.3612

Corrected Akaike information criterion (AICc) score: 1290144.9491

Bayesian information criterion (BIC) score: 1292675.6957

Total tree length (sum of branch lengths): 0.4046

Sum of internal branch lengths: 0.2724 (67.3318% of tree length)

WARNING: 49 near-zero internal branches (<0.0000) should be treated with caution

Such branches are denoted by '**' in the figure below

NOTE: Tree is UNROOTED although outgroup taxon 'U18466.2_African_swine_fever_virus_strain_BA71V_complete_genome' is drawn at root

+--U18466.2_African_swine_fever_virus_strain_BA71V_complete_genome

|

| +--KM262844.1_African_swine_fever_virus_strain_L60_complete_genome

| +--|

| | | +--KM102979.1_African_swine_fever_virus_isolate_26544/OG10_from_Italy_complete_genome

| | | +**|

| | | | | +--MN270979.1_African_swine_fever_virus_isolate_97/Ot/2012_complete_genome

| | | | | +--|

| | | | | | | +--MW736597.1_African_swine_fever_virus_strain_47039_complete_genome

| | | | | | | +**|

| | | | | | | | +--MW736600.1_African_swine_fever_virus_strain_30322_complete_genome

| | | | | | | +**|

| | | | | | | | +**MW736608.1_African_swine_fever_virus_strain_113049_WB_complete_genome

| | | | | | | +**|

| | | | | | | | +**MW736603.1_African_swine_fever_virus_strain_63525_WB_complete_genome

| | | | | | | +**|

| | | | | | | | | +--MW736605.1_African_swine_fever_virus_strain_51268_complete_genome

| | | | | | | | +**|

| | | | | | | | +--MW788408.1_African_swine_fever_virus_strain_35479_2014_genomic_sequence

| | | | | | | +**|

| | | | | | | | +**MW736598.1_African_swine_fever_virus_strain_2019_WB_complete_genome

| | | | | | | +**|

| | | | | | | | +--MW788407.1_African_swine_fever_virus_strain_31479_2005_genomic_sequence

| | | | | | +--|

| | | | | | | +--MW736601.1_African_swine_fever_virus_strain_49179_WB_complete_genome

| | | | | | +**|

| | | | | | | +--MW736604.1_African_swine_fever_virus_strain_15998_complete_genome

| | | | | | | +**|

| | | | | | | | +--MW736611.1_African_swine_fever_virus_strain_56140_genomic_sequence

| | | | | | +--|

| | | | | | +--MW736602.1_African_swine_fever_virus_strain_53706_genomic_sequence

| | | | +--|

| | | | | +**MW723492.1_African_swine_fever_virus_strain_4996_WB_complete_genome

| | | | +**|

| | | | +**MW788405.1_African_swine_fever_virus_strain_1537_WB_genomic_sequence

| | | +--|

| | | | | +**MN270980.1_African_swine_fever_virus_isolate_22653/Ca/2014_complete_genome

| | | | +--|

| | | | | +**MT932578.1_African_swine_fever_virus_strain_103917/18_partial_genome

| | | | +**|

| | | | | +**MT932579.1_African_swine_fever_virus_strain_55234/18_partial_genome

| | | | +--|

| | | | +--MW736606.1_African_swine_fever_virus_strain_34403_genomic_sequence

| | | +**|

| | | | | +--MW723494.1_African_swine_fever_virus_strain_23221_complete_genome

| | | | | +--|

| | | | | | +--MW723498.1_African_swine_fever_virus_strain_72912_WB_complete_genome

| | | | +--|

| | | | +**MW788406.1_African_swine_fever_virus_strain_22943_2008_genomic_sequence

| | | +--|

| | | | | +**KX354450.1_African_swine_fever_virus_isolate_47/Ss/2008_complete_genome

| | | | | +--|

| | | | | | +--MW723493.1_African_swine_fever_virus_strain_46830_complete_genome

| | | | | +**|

| | | | | | | +**MW723495.1_African_swine_fever_virus_strain_72398_WB_complete_genome

| | | | | | +**|

| | | | | | +**MW723499.1_African_swine_fever_virus_strain_22137_complete_genome

| | | | | +**|

| | | | | | +**MN270978.1_African_swine_fever_virus_isolate_72407/Ss/2005_complete_genome

| | | | | +--|

| | | | | | +--MW788410.1_African_swine_fever_virus_strain_25185_2008_genomic_sequence

| | | | | +**|

| | | | | | | +**MW723496.1_African_swine_fever_virus_strain_74377_complete_genome

| | | | | | +**|

| | | | | | +--MW723500.1_African_swine_fever_virus_strain_44076_complete_genome

| | | | | +--|

| | | | | | | +**MN270976.1_African_swine_fever_virus_isolate_60/Nu/1997_complete_genome

| | | | | | | +**|

| | | | | | | | | +**MN270977.1_African_swine_fever_virus_isolate_26/Ss/2004_complete_genome

| | | | | | | | +--|

| | | | | | | | | +--MW736599.1_African_swine_fever_virus_strain_98039_complete_genome

| | | | | | | | | +**|

| | | | | | | | | | +--MW736613.1_African_swine_fever_virus_strain_33747_WB_complete_genome

| | | | | | | | +--|

| | | | | | | | +--MW736612.1_African_swine_fever_virus_strain_31208_complete_genome

| | | | | | +--|

| | | | | | +--MW723497.1_African_swine_fever_virus_strain_22649_complete_genome

| | | | +**|

| | | | +--MW788411.1_UNVERIFIED__African_swine_fever_virus_strain_24225_2002_genomic_sequence

| | | +--|

| | | | | +**MW723484.1_African_swine_fever_virus_strain_Nu1991_2_complete_genome

| | | | | +--|

| | | | | | +--MW723486.1_African_swine_fever_virus_strain_Nu1991_7_complete_genome

| | | | +**|

| | | | +--MW723488.1_African_swine_fever_virus_strain_Nu1993_2_complete_genome

| | | +--|

| | | | | +--MN270973.1_African_swine_fever_virus_isolate_85/Ca/1985_complete_genome

| | | | | +--|

| | | | | | | +**MN270974.1_African_swine_fever_virus_isolate_141/Nu/1990_complete_genome

| | | | | | +--|

| | | | | | | +**MN270975.1_African_swine_fever_virus_isolate_142/Nu/1995_complete_genome

| | | | | | +--|

| | | | | | | +--MW723490.1_African_swine_fever_virus_strain_Nu1995_3_complete_genome

| | | | | | +--|

| | | | | | | +**MW723491.1_African_swine_fever_virus_strain_Nu1995_4_complete_genome

| | | | | | +**|

| | | | | | +**MW723489.1_African_swine_fever_virus_strain_Nu1995_2_complete_genome

| | | | +--|

| | | | | +**MW723483.1_African_swine_fever_virus_strain_Nu1990_1_complete_genome

| | | | +**|

| | | | | +--MW723485.1_African_swine_fever_virus_strain_Nu1991_3_complete_genome

| | | | +**|

| | | | +--MW723487.1_African_swine_fever_virus_strain_Or1993_1_complete_genome

| | | +--|

| | | | +--MW723482.1_African_swine_fever_virus_strain_Nu1986_complete_genome

| | | +--|

| | | | +**MN270972.1_African_swine_fever_virus_isolate_140/Or/1985_complete_genome

| | | +--|

| | | | | +**MN270969.1_African_swine_fever_virus_isolate_56/Ca/1978_complete_genome

| | | | | +**|

| | | | | | | +**MN270971.1_African_swine_fever_virus_isolate_139/Nu/1981_complete_genome

| | | | | | | +--|

| | | | | | | | +--MW800838.1_African_swine_fever_virus_strain_Or_1984_complete_genome

| | | | | | +--|

| | | | | | +**MW788409.1_UNVERIFIED__African_swine_fever_virus_strain_SS_1981_genomic_sequence

| | | | +**|

| | | | | +**MN270970.1_African_swine_fever_virus_isolate_57/Ca/1979_complete_genome

| | | | +**|

| | | | | +--MW723480.1_African_swine_fever_virus_strain_Ca1978_2_complete_genome

| | | | +**|

| | | | +**MW723481.1_African_swine_fever_virus_strain_Nu1979_complete_genome

| | | +--|

| | | | +--AM712239.1_African_swine_fever_virus_Benin_97/1_pathogenic_isolate_complete_genome

| | +--|

| | +--FN557520.1_African_swine_fever_virus_E75_complete_genome_strain_E75

+--|

| | +--AY261360.1_African_swine_fever_virus_isolate_Kenya_1950_complete_genome

| | +----|

| | | | +--MT956648.1_African_swine_fever_virus_isolate_Uvira_B53_complete_genome

| | | +--|

| | | +--MW856067.1_African_swine_fever_virus_strain_BUR/18/Rutana_complete_genome

| | +--------------------------------------|

| | | | +--KM111295.1_African_swine_fever_virus_strain_Ken06.Bus_complete_genome

| | | +------|

| | | | +**MH025916.1_African_swine_fever_virus_strain_R8_complete_genome

| | | +--|

| | | +**MH025917.1_African_swine_fever_virus_strain_R7_complete_genome

| | +--|

| | | | +--AY261366.1_African_swine_fever_virus_isolate_Warthog_complete_genome

| | | | +-----|

| | | | | | +-------------MN318203.3_African_swine_fever_virus_isolate_LIV_5_40_complete_genome

| | | | | | +--|

| | | | | | | | +---MN336500.3_African_swine_fever_virus_isolate_RSA_2_2008_complete_genome

| | | | | | | +--|

| | | | | | | +--MN394630.3_African_swine_fever_virus_isolate_SPEC_57_complete_genome

| | | | | | +--|

| | | | | | | | +--MN641877.2_African_swine_fever_virus_isolate_RSA_2_2004_complete_genome

| | | | | | | +--|

| | | | | | | +--MN630494.2_African_swine_fever_virus_isolate_Zaire_complete_genome

| | | | | +--------------|

| | | | | +--MN641876.2_African_swine_fever_virus_isolate_RSA_W1_1999_complete_genome

| | | +--|

| | | +-------AY261364.1_African_swine_fever_virus_isolate_Tengani_62_complete_genome

| | +--|

| | | | +**FR682468.2_African_swine_fever_virus_isolate_ASFV_Georgia_2007/1_genome_assembly_complete_genome__monopartite

| | | | +**|

| | | | | | +**MG939583.1_UNVERIFIED__African_swine_fever_virus_isolate_Pol16_20186_o7_complete_genome

| | | | | | +--|

| | | | | | | +--MG939584.1_UNVERIFIED__African_swine_fever_virus_isolate_Pol16_20538_o9_complete_genome

| | | | | | +**|

| | | | | | | +--MN194591.1_African_swine_fever_virus_isolate_ASFV/Kyiv/2016/131_complete_genome

| | | | | | +--|

| | | | | | | +--MT847621.1_African_swine_fever_virus_isolate_Pol18_28298_O111_complete_genome

| | | | | | +**|

| | | | | | | +--MG939588.1_UNVERIFIED__African_swine_fever_virus_isolate_Pol17_04461_C210_complete_genome

| | | | | | +**|

| | | | | | | +--MT847622.1_African_swine_fever_virus_isolate_Pol17_31177_O81_complete_genome

| | | | | | +--|

| | | | | | | | +**MT847620.1_African_swine_fever_virus_isolate_Pol17_55892_C754_complete_genome

| | | | | | | +--|

| | | | | | | +**MT847623.2_African_swine_fever_virus_isolate_Pol19_53050_C1959/19_complete_genome

| | | | | +--|

| | | | | | +--MH766894.2_UNVERIFIED__African_swine_fever_virus_isolate_ASFV-SY18_complete_genome

| | | | | | +--|

| | | | | | | | +**MT166692.1_African_swine_fever_virus_strain_ASFV_Hanoi_2019_partial_genome

| | | | | | | | +---|

| | | | | | | | | +--LS478113.1_African_swine_fever_virus_isolate_Estonia_2014_genome_assembly_complete_genome__monopartite

| | | | | | | +--|

| | | | | | | +--MW656282.1_African_swine_fever_virus_isolate_Pig/Heilongjiang/HRB1/2020_complete_genome

| | | | | | +**|

| | | | | | | | +--233_LILO_scaffold

| | | | | | | | +**|

| | | | | | | | | +--ASF-10_LILO_scaffold

| | | | | | | | +--|

| | | | | | | | | +--ASF-30_LILO_scaffold

| | | | | | | +**|

| | | | | | | +--3142_LILO_scaffold

| | | | | | +**|

| | | | | | | | +**MN393476.1_African_swine_fever_virus_isolate_ASFV_Wuhan_2019-1_complete_genome

| | | | | | | +--|

| | | | | | | +**MN393477.1_African_swine_fever_virus_isolate_ASFV_Wuhan_2019-2_complete_genome

| | | | | | +**|

| | | | | | | +--MW396979.1_African_swine_fever_virus_isolate_ASFV/Timor-Leste/2019/1_complete_genome

| | | | | | +**|

| | | | | | | +--MW306191.1_African_swine_fever_virus_isolate_ASFV/Primorsky_19/WB-6723_complete_genome

| | | | | | +**|

| | | | | | | +--MT496893.1_African_swine_fever_virus_isolate_GZ201801_complete_genome

| | | | | | +**|

| | | | | | | +--MN715134.1_African_swine_fever_virus_strain_ASFV_HU_2018_complete_genome

| | | | | | +**|

| | | | | | | +--MW306190.1_African_swine_fever_virus_isolate_ASFV/Amur_19/WB-6905_complete_genome

| | | | | | +**|

| | | | | | | +**MT748042.1_UNVERIFIED__African_swine_fever_virus_strain_ASFV/Korea/pig/PaJu1/2019_complete_genome

| | | | | | +**|

| | | | | | | +--MN172368.1_African_swine_fever_virus_strain_ASFV/pig/China/CAS19-01/2019_complete_genome

| | | | | | +**|

| | | | | | | | +--MK645909.1_African_swine_fever_virus_isolate_ASFV-wbBS01_complete_genome

| | | | | | | +--|

| | | | | | | +--MK940252.1_African_swine_fever_virus_isolate_CN/2019/InnerMongolia-AES01_complete_genome

| | | | | | +**|

| | | | | | | +--MT180393.1_African_swine_fever_virus_strain_ASFV_NgheAn_2019_partial_genome

| | | | | | +--|

| | | | | | | | +--126_LILO_scaffold

| | | | | | | +--|

| | | | | | | | +--237_LILO_scaffold

| | | | | | | | +--|

| | | | | | | | | +--2607_LILO_scaffold

| | | | | | | +**|

| | | | | | | +--261_LILO_scaffold

| | | | | +**|

| | | | | +**MK543947.1_African_swine_fever_virus_strain_Belgium/Etalle/wb/2018_complete_genome

| | | | +--|

| | | | | | +--KJ747406.1_UNVERIFIED__African_swine_fever_virus_isolate_Kashino_04/13_genomic_sequence

| | | | | | +--|

| | | | | | | | +--MK628478.1_African_swine_fever_virus_isolate_ASFV/LT14/1490_complete_genome

| | | | | | | +--|

| | | | | | | +--MW306192.1_African_swine_fever_virus_isolate_ASFV/Ulyanovsk_19/WB-5699_complete_genome

| | | | | +**|

| | | | | +--MT459800.1_African_swine_fever_virus_isolate_ASFV/Kabardino-Balkaria_19/WB-964_complete_genome

| | | | +--|

| | | | | +--MW856068.1_African_swine_fever_virus_strain_MAL/19/Karonga_complete_genome

| | | +--|

| | | +--OK236383.1_UNVERIFIED__African_swine_fever_virus_isolate_ASF/IND/20/CAD/543_partial_genome

| | +--|

| | | +--MN913970.1_African_swine_fever_virus_strain_Liv13/33__OmLF2__complete_genome

| +--|

| +--MZ202520.1_African_swine_fever_virus_strain_K49_complete_genome

|

+--AM712240.1_African_swine_fever_virus_OURT_88/3__avirulent_field_isolate__complete_genome

Tree in newick format:

(U18466.2_African_swine_fever_virus_strain_BA71V_complete_genome:0.0030841644,((KM262844.1_African_swine_fever_virus_strain_L60_complete_genome:0.0000864738,(((((((((((KM102979.1_African_swine_fever_virus_isolate_26544/OG10_from_Italy_complete_genome:0.0000398664,((MN270979.1_African_swine_fever_virus_isolate_97/Ot/2012_complete_genome:0.0000110385,(((((((MW736597.1_African_swine_fever_virus_strain_47039_complete_genome:0.0000110356,MW736600.1_African_swine_fever_virus_strain_30322_complete_genome:0.0000054515):0.0000004793,MW736608.1_African_swine_fever_virus_strain_113049_WB_complete_genome:0.0000004793):0.0000004793,MW736603.1_African_swine_fever_virus_strain_63525_WB_complete_genome:0.0000004793):0.0000004793,(MW736605.1_African_swine_fever_virus_strain_51268_complete_genome:0.0000170869,MW788408.1_African_swine_fever_virus_strain_35479_2014_genomic_sequence:0.0000054633):0.0000004793):0.0000004793,MW736598.1_African_swine_fever_virus_strain_2019_WB_complete_genome:0.0000004793):0.0000004793,MW788407.1_African_swine_fever_virus_strain_31479_2005_genomic_sequence:0.0000112181):0.0000004793,(MW736601.1_African_swine_fever_virus_strain_49179_WB_complete_genome:0.0000054502,((MW736604.1_African_swine_fever_virus_strain_15998_complete_genome:0.0000110356,MW736611.1_African_swine_fever_virus_strain_56140_genomic_sequence:0.0000110356):0.0000004793,MW736602.1_African_swine_fever_virus_strain_53706_genomic_sequence:0.0000110323):0.0000110321):0.0000004793):0.0000054495):0.0000110309,(MW723492.1_African_swine_fever_virus_strain_4996_WB_complete_genome:0.0000004793,MW788405.1_African_swine_fever_virus_strain_1537_WB_genomic_sequence:0.0000004793):0.0000004793):0.0000054493):0.0000004793,(MN270980.1_African_swine_fever_virus_isolate_22653/Ca/2014_complete_genome:0.0000004793,(MT932578.1_African_swine_fever_virus_strain_103917/18_partial_genome:0.0000004793,(MT932579.1_African_swine_fever_virus_strain_55234/18_partial_genome:0.0000004793,MW736606.1_African_swine_fever_virus_strain_34403_genomic_sequence:0.0000054531):0.0000170866):0.0000004793):0.0000398629):0.0000110293,((MW723494.1_African_swine_fever_virus_strain_23221_complete_genome:0.0000057081,MW723498.1_African_swine_fever_virus_strain_72912_WB_complete_genome:0.0000283926):0.0000053372,MW788406.1_African_swine_fever_virus_strain_22943_2008_genomic_sequence:0.0000013156):0.0000398015):0.0000004793,(((((((KX354450.1_African_swine_fever_virus_isolate_47/Ss/2008_complete_genome:0.0000004793,MW723493.1_African_swine_fever_virus_strain_46830_complete_genome:0.0000054532):0.0000341746,(MW723495.1_African_swine_fever_virus_strain_72398_WB_complete_genome:0.0000004793,MW723499.1_African_swine_fever_virus_strain_22137_complete_genome:0.0000004793):0.0000004793):0.0000004793,MN270978.1_African_swine_fever_virus_isolate_72407/Ss/2005_complete_genome:0.0000004793):0.0000004793,MW788410.1_African_swine_fever_virus_strain_25185_2008_genomic_sequence:0.0000054607):0.0000054508,(MW723496.1_African_swine_fever_virus_strain_74377_complete_genome:0.0000004793,MW723500.1_African_swine_fever_virus_strain_44076_complete_genome:0.0000110343):0.0000004793):0.0000004793,((MN270976.1_African_swine_fever_virus_isolate_60/Nu/1997_complete_genome:0.0000004793,(MN270977.1_African_swine_fever_virus_isolate_26/Ss/2004_complete_genome:0.0000004793,((MW736599.1_African_swine_fever_virus_strain_98039_complete_genome:0.0000055225,MW736613.1_African_swine_fever_virus_strain_33747_WB_complete_genome:0.0000110353):0.0000004793,MW736612.1_African_swine_fever_virus_strain_31208_complete_genome:0.0000054531):0.0000110344):0.0000054508):0.0000004793,MW723497.1_African_swine_fever_virus_strain_22649_complete_genome:0.0000054514):0.0000170838):0.0000110330,MW788411.1_UNVERIFIED__African_swine_fever_virus_strain_24225_2002_genomic_sequence:0.0000059324):0.0000004793):0.0000569407,((MW723484.1_African_swine_fever_virus_strain_Nu1991_2_complete_genome:0.0000004793,MW723486.1_African_swine_fever_virus_strain_Nu1991_7_complete_genome:0.0000054530):0.0000110350,MW723488.1_African_swine_fever_virus_strain_Nu1993_2_complete_genome:0.0000110349):0.0000004793):0.0000054494,((MN270973.1_African_swine_fever_virus_isolate_85/Ca/1985_complete_genome:0.0000170888,(MN270974.1_African_swine_fever_virus_isolate_141/Nu/1990_complete_genome:0.0000004793,(MN270975.1_African_swine_fever_virus_isolate_142/Nu/1995_complete_genome:0.0000004793,(MW723490.1_African_swine_fever_virus_strain_Nu1995_3_complete_genome:0.0000054505,(MW723491.1_African_swine_fever_virus_strain_Nu1995_4_complete_genome:0.0000004793,MW723489.1_African_swine_fever_virus_strain_Nu1995_2_complete_genome:0.0000004793):0.0000004793):0.0000054501):0.0000170783):0.0000054516):0.0000054156,(MW723483.1_African_swine_fever_virus_strain_Nu1990_1_complete_genome:0.0000004793,(MW723485.1_African_swine_fever_virus_strain_Nu1991_3_complete_genome:0.0000110376,MW723487.1_African_swine_fever_virus_strain_Or1993_1_complete_genome:0.0000473726):0.0000004793):0.0000004793):0.0000270408):0.0000881630,MW723482.1_African_swine_fever_virus_strain_Nu1986_complete_genome:0.0000170788):0.0000202099,MN270972.1_African_swine_fever_virus_isolate_140/Or/1985_complete_genome:0.0000004793):0.0000284047,((MN270969.1_African_swine_fever_virus_isolate_56/Ca/1978_complete_genome:0.0000004793,((MN270971.1_African_swine_fever_virus_isolate_139/Nu/1981_complete_genome:0.0000004793,MW800838.1_African_swine_fever_virus_strain_Or_1984_complete_genome:0.0000341018):0.0000054429,MW788409.1_UNVERIFIED__African_swine_fever_virus_strain_SS_1981_genomic_sequence:0.0000004793):0.0000054425):0.0000004793,(MN270970.1_African_swine_fever_virus_isolate_57/Ca/1979_complete_genome:0.0000004793,(MW723480.1_African_swine_fever_virus_strain_Ca1978_2_complete_genome:0.0000054424,MW723481.1_African_swine_fever_virus_strain_Nu1979_complete_genome:0.0000004793):0.0000004793):0.0000004793):0.0000004793):0.0000397949,AM712239.1_African_swine_fever_virus_Benin_97/1_pathogenic_isolate_complete_genome:0.0002513162):0.0000287063,FN557520.1_African_swine_fever_virus_E75_complete_genome_strain_E75:0.0000815374):0.0000581433):0.0000297956,((((((AY261360.1_African_swine_fever_virus_isolate_Kenya_1950_complete_genome:0.0066955848,(MT956648.1_African_swine_fever_virus_isolate_Uvira_B53_complete_genome:0.0007391911,MW856067.1_African_swine_fever_virus_strain_BUR/18/Rutana_complete_genome:0.0013051426):0.0077031001):0.0131690045,(KM111295.1_African_swine_fever_virus_strain_Ken06.Bus_complete_genome:0.0003868076,(MH025916.1_African_swine_fever_virus_strain_R8_complete_genome:0.0000004793,MH025917.1_African_swine_fever_virus_strain_R7_complete_genome:0.0000004793):0.0005643651):0.0169441403):0.0950134940,((AY261366.1_African_swine_fever_virus_isolate_Warthog_complete_genome:0.0000538874,(((MN318203.3_African_swine_fever_virus_isolate_LIV_5_40_complete_genome:0.0341678090,(MN336500.3_African_swine_fever_virus_isolate_RSA_2_2008_complete_genome:0.0098959593,MN394630.3_African_swine_fever_virus_isolate_SPEC_57_complete_genome:0.0080636826):0.0061922733):0.0052435199,(MN641877.2_African_swine_fever_virus_isolate_RSA_2_2004_complete_genome:0.0094610066,MN630494.2_African_swine_fever_virus_isolate_Zaire_complete_genome:0.0066731917):0.0079258669):0.0087521139,MN641876.2_African_swine_fever_virus_isolate_RSA_W1_1999_complete_genome:0.0046996372):0.0378077967):0.0165616208,AY261364.1_African_swine_fever_virus_isolate_Tengani_62_complete_genome:0.0196027649):0.0039306088):0.0076002502,((((FR682468.2_African_swine_fever_virus_isolate_ASFV_Georgia_2007/1_genome_assembly_complete_genome__monopartite:0.0000004793,(((((((MG939583.1_UNVERIFIED__African_swine_fever_virus_isolate_Pol16_20186_o7_complete_genome:0.0000004793,MG939584.1_UNVERIFIED__African_swine_fever_virus_isolate_Pol16_20538_o9_complete_genome:0.0001141324):0.0000104003,MN194591.1_African_swine_fever_virus_isolate_ASFV/Kyiv/2016/131_complete_genome:0.0000825480):0.0000004793,MT847621.1_African_swine_fever_virus_isolate_Pol18_28298_O111_complete_genome:0.0000053340):0.0000058941,MG939588.1_UNVERIFIED__African_swine_fever_virus_isolate_Pol17_04461_C210_complete_genome:0.0000053345):0.0000004793,MT847622.1_African_swine_fever_virus_isolate_Pol17_31177_O81_complete_genome:0.0000053338):0.0000004793,(MT847620.1_African_swine_fever_virus_isolate_Pol17_55892_C754_complete_genome:0.0000004793,MT847623.2_African_swine_fever_virus_isolate_Pol19_53050_C1959/19_complete_genome:0.0000004793):0.0000053338):0.0000192177,((((((((((((((MH766894.2_UNVERIFIED__African_swine_fever_virus_isolate_ASFV-SY18_complete_genome:0.0001331790,((MT166692.1_African_swine_fever_virus_strain_ASFV_Hanoi_2019_partial_genome:0.0000019816,LS478113.1_African_swine_fever_virus_isolate_Estonia_2014_genome_assembly_complete_genome__monopartite:0.0006299677):0.0105968980,MW656282.1_African_swine_fever_virus_isolate_Pig/Heilongjiang/HRB1/2020_complete_genome:0.0001877649):0.0000248272):0.0000050521,(((233_LILO_scaffold:0.0000386718,ASF-10_LILO_scaffold:0.0000592247):0.0000004793,ASF-30_LILO_scaffold:0.0000112412):0.0000080050,3142_LILO_scaffold:0.0000674998):0.0000004793):0.0000004793,(MN393476.1_African_swine_fever_virus_isolate_ASFV_Wuhan_2019-1_complete_genome:0.0000004793,MN393477.1_African_swine_fever_virus_isolate_ASFV_Wuhan_2019-2_complete_genome:0.0000004793):0.0000104039):0.0000004793,MW396979.1_African_swine_fever_virus_isolate_ASFV/Timor-Leste/2019/1_complete_genome:0.0000053344):0.0000011160,MW306191.1_African_swine_fever_virus_isolate_ASFV/Primorsky_19/WB-6723_complete_genome:0.0000269842):0.0000009877,MT496893.1_African_swine_fever_virus_isolate_GZ201801_complete_genome:0.0000815531):0.0000004793,MN715134.1_African_swine_fever_virus_strain_ASFV_HU_2018_complete_genome:0.0000103964):0.0000004793,MW306190.1_African_swine_fever_virus_isolate_ASFV/Amur_19/WB-6905_complete_genome:0.0000269848):0.0000004793,MT748042.1_UNVERIFIED__African_swine_fever_virus_strain_ASFV/Korea/pig/PaJu1/2019_complete_genome:0.0000004793):0.0000004793,MN172368.1_African_swine_fever_virus_strain_ASFV/pig/China/CAS19-01/2019_complete_genome:0.0000053341):0.0000004793,(MK645909.1_African_swine_fever_virus_isolate_ASFV-wbBS01_complete_genome:0.0000485710,MK940252.1_African_swine_fever_virus_isolate_CN/2019/InnerMongolia-AES01_complete_genome:0.0000706767):0.0000053319):0.0000004793,MT180393.1_African_swine_fever_virus_strain_ASFV_NgheAn_2019_partial_genome:0.0000053769):0.0000012465,(126_LILO_scaffold:0.0000429481,((237_LILO_scaffold:0.0000746450,2607_LILO_scaffold:0.0000124326):0.0000058092,261_LILO_scaffold:0.0000415538):0.0000010184):0.0000067929):0.0000068803,MK543947.1_African_swine_fever_virus_strain_Belgium/Etalle/wb/2018_complete_genome:0.0000004793):0.0000012597):0.0000228474):0.0000010184,((KJ747406.1_UNVERIFIED__African_swine_fever_virus_isolate_Kashino_04/13_genomic_sequence:0.0001135993,(MK628478.1_African_swine_fever_virus_isolate_ASFV/LT14/1490_complete_genome:0.0000377775,MW306192.1_African_swine_fever_virus_isolate_ASFV/Ulyanovsk_19/WB-5699_complete_genome:0.0000869079):0.0000161872):0.0000058611,MT459800.1_African_swine_fever_virus_isolate_ASFV/Kabardino-Balkaria_19/WB-964_complete_genome:0.0000540144):0.0000004793):0.0000357602,MW856068.1_African_swine_fever_virus_strain_MAL/19/Karonga_complete_genome:0.0003786708):0.0052370620,OK236383.1_UNVERIFIED__African_swine_fever_virus_isolate_ASF/IND/20/CAD/543_partial_genome:0.0042656094):0.0092474776):0.0070544071,MN913970.1_African_swine_fever_virus_strain_Liv13/33__OmLF2__complete_genome:0.0090757834):0.0073000285,MZ202520.1_African_swine_fever_virus_strain_K49_complete_genome:0.0076995305):0.0042310228):0.0010187151,AM712240.1_African_swine_fever_virus_OURT_88/3__avirulent_field_isolate__complete_genome:0.0024203482);
