## Supplementary for "No part gets left behind: Tiled nanopore sequencing of whole ASFV genomes stitched together using Lilo": Supplementary Document S2_ tree details for 5B.docx

Supplementary document X- Output from iqtree2 detailing model selection and full likelihood statistics for the tree in figure 5B.

IQ-TREE 2.0.5 built May 15 2020

Type of analysis: ModelFinder + tree reconstruction

Random seed number: 938260

REFERENCES

----------

To cite IQ-TREE please use:

Bui Quang Minh, Heiko A. Schmidt, Olga Chernomor, Dominik Schrempf,

Michael D. Woodhams, Arndt von Haeseler, and Robert Lanfear (2020)

IQ-TREE 2: New models and efficient methods for phylogenetic inference

in the genomic era. Mol. Biol. Evol., in press.

https://doi.org/10.1093/molbev/msaa015

To cite ModelFinder please use:

Subha Kalyaanamoorthy, Bui Quang Minh, Thomas KF Wong, Arndt von Haeseler,

and Lars S Jermiin (2017) ModelFinder: Fast model selection for

accurate phylogenetic estimates. Nature Methods, 14:587–589.

https://doi.org/10.1038/nmeth.4285

SEQUENCE ALIGNMENT

------------------

Input data: 36 sequences with 188016 nucleotide sites

Number of constant sites: 187735 (= 99.8505% of all sites)

Number of invariant (constant or ambiguous constant) sites: 187735 (= 99.8505% of all sites)

Number of parsimony informative sites: 27

Number of distinct site patterns: 470

ModelFinder

-----------

Best-fit model according to BIC: HKY+F+I

List of models sorted by BIC scores:

Model	LogL	AIC	w-AIC	AICc	w-AICc	BIC	w-BIC

HKY+F+I	-258342.353 516832.707 - 0.00373 516832.766 - 0.00374 517583.384 +	0.755

TPM3+F+I	-258338.300 516826.599 +	0.0791 516826.660 +	0.0792 517587.421 +	0.1

TPM3u+F+I	-258338.304 516826.607 +	0.0788 516826.668 +	0.0789 517587.428 +	0.0999

TPM2u+F+I	-258339.935 516829.869 -	0.0154 516829.930 -	0.0154 517590.690 -	0.0196

TPM2+F+I	-258339.935 516829.869 -	0.0154 516829.930 -	0.0154 517590.690 -	0.0196

TN+F+I	-258341.873 516833.745 - 0.00222 516833.806 - 0.00222 517594.566 - 0.00282

K3Pu+F+I	-258341.988 516833.976 - 0.00198 516834.037 - 0.00198 517594.798 - 0.00251

TIM3+F+I	-258337.823 516827.647 -	0.0468 516827.709 -	0.0469 517598.612 - 0.000372

TIM2+F+I	-258339.800 516831.600 - 0.00649 516831.662 -	0.0065 517602.565 - 5.16e-05

TVM+F+I	-258334.525 516823.050 +	0.467 516823.114 +	0.466 517604.160 - 2.33e-05

TIM+F+I	-258341.484 516834.967 - 0.00121 516835.029 - 0.00121 517605.933 - 9.58e-06

GTR+F+I	-258334.028 516824.055 +	0.282 516824.121 +	0.282 517615.309 - 8.82e-08

GTR+F+G4	-258345.823 516847.647 - 2.13e-06 516847.713 - 2.12e-06 517638.901 - 6.64e-13

GTR+F+I+G4	-258341.012 516840.024 - 9.61e-05 516840.091 - 9.6e-05 517641.423 - 1.88e-13

GTR+F	-258359.695 516873.390 - 5.47e-12 516873.454 - 5.47e-12 517654.500 - 2.72e-16

GTR+F+R2	-258353.324 516864.649 - 4.32e-10 516864.716 - 4.32e-10 517666.047 - 8.47e-19

GTR+F+R3	-258350.550 516863.099 - 9.38e-10 516863.170 - 9.35e-10 517684.786 - 7.22e-23

F81+F+I	-258419.180 516984.360 - 4.37e-36 516984.418 - 4.39e-36 517724.893 - 1.41e-31

K2P+I	-263458.137 527058.273 -	0 527058.328 -	0 527778.517 -	0

TVMe+I	-263443.170 527034.341 -	0 527034.400 -	0 527785.018 -	0

TIM3e+I	-263450.224 527046.447 -	0 527046.504 -	0 527786.980 -	0

K3P+I	-263457.226 527058.452 -	0 527058.508 -	0 527788.841 -	0

TNe+I	-263457.889 527059.779 -	0 527059.835 -	0 527790.167 -	0

TIM2e+I	-263452.404 527050.809 -	0 527050.866 -	0 527791.341 -	0

SYM+I	-263442.374 527034.748 -	0 527034.809 -	0 527795.570 -	0

TIMe+I	-263456.727 527059.454 -	0 527059.511 -	0 527799.986 -	0

JC+I	-263529.744 527199.488 -	0 527199.541 -	0 527909.588 -	0

AIC, w-AIC	: Akaike information criterion scores and weights.

AICc, w-AICc : Corrected AIC scores and weights.

BIC, w-BIC	: Bayesian information criterion scores and weights.

Plus signs denote the 95% confidence sets.

Minus signs denote significant exclusion.

SUBSTITUTION PROCESS

--------------------

Model of substitution: HKY+F+I

Rate parameter R:

A-C: 1.0000

A-G: 4.4564

A-T: 1.0000

C-G: 1.0000

C-T: 4.4564

G-T: 1.0000

State frequencies: (empirical counts from alignment)

pi(A) = 0.307

pi(C) = 0.1927

pi(G) = 0.192

pi(T) = 0.3083

Rate matrix Q:

A	-0.8725	0.124	0.5503	0.1983

C	0.1974	-1.204	0.1235	0.8835

G	0.8798	0.124	-1.202	0.1983

T	0.1974	0.5524	0.1235	-0.8733

Model of rate heterogeneity: Invar

Proportion of invariable sites: 0.9755

Category Relative_rate Proportion

0	0	0.9755

1	40.82	0.0245

MULTIPLE RUNS

-------------

Run	logL

1	-258340.3878

2	-258340.3893

3	-258340.2488

4	-258340.4544

5	-258340.4244

6	-258340.3858

7	-258340.3847

8	-258340.3829

9	-258340.4275

10	-258340.3550

11	-258340.3499

12	-258340.3914

13	-258340.3817

14	-258340.3667

15	-258340.3805

16	-258340.3903

17	-258340.3928

18	-258340.3497

19	-258340.3604

20	-258340.4505

MAXIMUM LIKELIHOOD TREE

-----------------------

Log-likelihood of the tree: -258340.2488 (s.e. 8591427046549753119504488702157545671920426418176.0000)

Unconstrained log-likelihood (without tree): -432326.6493

Number of free parameters (#branches + #model parameters): 74

Akaike information criterion (AIC) score: 516828.4976

Corrected Akaike information criterion (AICc) score: 516828.5567

Bayesian information criterion (BIC) score: 517579.1745

Total tree length (sum of branch lengths): 0.0016

Sum of internal branch lengths: 0.0001 (7.5090% of tree length)

WARNING: 21 near-zero internal branches (<0.0000) should be treated with caution

Such branches are denoted by '**' in the figure below

NOTE: Tree is UNROOTED although outgroup taxon 'FR682468.2_African_swine_fever_virus_isolate_ASFV_Georgia_2007/1_genome_assembly_complete_genome__monopartite' is drawn at root

+**FR682468.2_African_swine_fever_virus_isolate_ASFV_Georgia_2007/1_genome_assembly_complete_genome__monopartite

|

| +--MG939583.1_UNVERIFIED__African_swine_fever_virus_isolate_Pol16_20186_o7_complete_genome

| +--|

| | | +-------------------MN194591.1_African_swine_fever_virus_isolate_ASFV/Kyiv/2016/131_complete_genome

| | +**|

| | +--MT847621.1_African_swine_fever_virus_isolate_Pol18_28298_O111_complete_genome

| +**|

| | | +**MT847620.1_African_swine_fever_virus_isolate_Pol17_55892_C754_complete_genome

| | +--|

| | +**MT847623.2_African_swine_fever_virus_isolate_Pol19_53050_C1959/19_complete_genome

| +**|

| | +--MT847622.1_African_swine_fever_virus_isolate_Pol17_31177_O81_complete_genome

| +---|

| | +--MG939588.1_UNVERIFIED__African_swine_fever_virus_isolate_Pol17_04461_C210_complete_genome

+----|

| | +------------------------------MH766894.2_UNVERIFIED__African_swine_fever_virus_isolate_ASFV-SY18_complete_genome

| | +**|

| | | +-------------------------------------------------MW656282.1_African_swine_fever_virus_isolate_Pig/Heilongjiang/HRB1/2020_complete_genome

| | +**|

| | | | +--MW396979.1_African_swine_fever_virus_isolate_ASFV/Timor-Leste/2019/1_complete_genome

| | | +**|

| | | | +-------PHL_233_LILO

| | | | +**|

| | | | | +------------PHL_10_LILO

| | | | +--|

| | | | | +--PHL_30_LILO

| | | +**|

| | | +---------------PHL_3142_LILO

| | +**|

| | | +-----MW306190.1_African_swine_fever_virus_isolate_ASFV/Amur_19/WB-6905_complete_genome

| | +**|

| | | +--MT180393.1_African_swine_fever_virus_strain_ASFV_NgheAn_2019_partial_genome

| | +**|

| | | +--MN715134.1_African_swine_fever_virus_strain_ASFV_HU_2018_complete_genome

| | +**|

| | | | +**MN393476.1_African_swine_fever_virus_isolate_ASFV_Wuhan_2019-1_complete_genome

| | | +--|

| | | +**MN393477.1_African_swine_fever_virus_isolate_ASFV_Wuhan_2019-2_complete_genome

| | +**|

| | | | +**MN172368.1_African_swine_fever_virus_strain_ASFV/pig/China/CAS19-01/2019_complete_genome

| | | +**|

| | | +-------------------MT166692.1_African_swine_fever_virus_strain_ASFV_Hanoi_2019_partial_genome

| | +**|

| | | +-----MW306191.1_African_swine_fever_virus_isolate_ASFV/Primorsky_19/WB-6723_complete_genome

| | +**|

| | | +**MT748042.1_UNVERIFIED__African_swine_fever_virus_strain_ASFV/Korea/pig/PaJu1/2019_complete_genome

| | +**|

| | | +------------------MT496893.1_African_swine_fever_virus_isolate_GZ201801_complete_genome

| | +**|

| | | | +----------MK645909.1_African_swine_fever_virus_isolate_ASFV-wbBS01_complete_genome

| | | +--|

| | | +---------------MK940252.1_African_swine_fever_virus_isolate_CN/2019/InnerMongolia-AES01_complete_genome

| | +--|

| | | | +---------PHL_126_LILO

| | | +--|

| | | | +----------------PHL_237_LILO

| | | | +--|

| | | | | +--PHL_2607_LILO

| | | +**|

| | | +--------PHL_261_LILO

| +**|

| +**MK543947.1_African_swine_fever_virus_strain_Belgium/Etalle/wb/2018_complete_genome

|

| +-------------------------KJ747406.1_UNVERIFIED__African_swine_fever_virus_isolate_Kashino_04/13_genomic_sequence

| +--|

| | | +-------MK628478.1_African_swine_fever_virus_isolate_ASFV/LT14/1490_complete_genome

| | +--|

| | +-------------------MW306192.1_African_swine_fever_virus_isolate_ASFV/Ulyanovsk_19/WB-5699_complete_genome

+**|

+-----------MT459800.1_African_swine_fever_virus_isolate_ASFV/Kabardino-Balkaria_19/WB-964_complete_genome

Tree in newick format:

(FR682468.2_African_swine_fever_virus_isolate_ASFV_Georgia_2007/1_genome_assembly_complete_genome__monopartite:0.0000005319,(((((MG939583.1_UNVERIFIED__African_swine_fever_virus_isolate_Pol16_20186_o7_complete_genome:0.0000106375,(MN194591.1_African_swine_fever_virus_isolate_ASFV/Kyiv/2016/131_complete_genome:0.0000840577,MT847621.1_African_swine_fever_virus_isolate_Pol18_28298_O111_complete_genome:0.0000053187):0.0000005319):0.0000063414,(MT847620.1_African_swine_fever_virus_isolate_Pol17_55892_C754_complete_genome:0.0000005319,MT847623.2_African_swine_fever_virus_isolate_Pol19_53050_C1959/19_complete_genome:0.0000005319):0.0000053187):0.0000005319,MT847622.1_African_swine_fever_virus_isolate_Pol17_31177_O81_complete_genome:0.0000053187):0.0000005319,MG939588.1_UNVERIFIED__African_swine_fever_virus_isolate_Pol17_04461_C210_complete_genome:0.0000053187):0.0000197725,(((((((((((((MH766894.2_UNVERIFIED__African_swine_fever_virus_isolate_ASFV-SY18_complete_genome:0.0001303401,MW656282.1_African_swine_fever_virus_isolate_Pig/Heilongjiang/HRB1/2020_complete_genome:0.0002106396):0.0000042740,(MW396979.1_African_swine_fever_virus_isolate_ASFV/Timor-Leste/2019/1_complete_genome:0.0000053187,(((PHL_233_LILO:0.0000358985,PHL_10_LILO:0.0000562338):0.0000005319,PHL_30_LILO:0.0000106375):0.0000080187,PHL_3142_LILO:0.0000666422):0.0000005319):0.0000005319):0.0000005319,MW306190.1_African_swine_fever_virus_isolate_ASFV/Amur_19/WB-6905_complete_genome:0.0000265944):0.0000005319,MT180393.1_African_swine_fever_virus_strain_ASFV_NgheAn_2019_partial_genome:0.0000053187):0.0000005319,MN715134.1_African_swine_fever_virus_strain_ASFV_HU_2018_complete_genome:0.0000106375):0.0000005319,(MN393476.1_African_swine_fever_virus_isolate_ASFV_Wuhan_2019-1_complete_genome:0.0000005319,MN393477.1_African_swine_fever_virus_isolate_ASFV_Wuhan_2019-2_complete_genome:0.0000005319):0.0000106375):0.0000005319,(MN172368.1_African_swine_fever_virus_strain_ASFV/pig/China/CAS19-01/2019_complete_genome:0.0000039366,MT166692.1_African_swine_fever_virus_strain_ASFV_Hanoi_2019_partial_genome:0.0000847071):0.0000011635):0.0000005319,MW306191.1_African_swine_fever_virus_isolate_ASFV/Primorsky_19/WB-6723_complete_genome:0.0000265944):0.0000005319,MT748042.1_UNVERIFIED__African_swine_fever_virus_strain_ASFV/Korea/pig/PaJu1/2019_complete_genome:0.0000005319):0.0000005319,MT496893.1_African_swine_fever_virus_isolate_GZ201801_complete_genome:0.0000803483):0.0000005319,(MK645909.1_African_swine_fever_virus_isolate_ASFV-wbBS01_complete_genome:0.0000478713,MK940252.1_African_swine_fever_virus_isolate_CN/2019/InnerMongolia-AES01_complete_genome:0.0000697315):0.0000053187):0.0000005319,(PHL_126_LILO:0.0000421084,((PHL_237_LILO:0.0000717177,PHL_2607_LILO:0.0000119818):0.0000057573,PHL_261_LILO:0.0000413702):0.0000005319):0.0000067998):0.0000066428,MK543947.1_African_swine_fever_virus_strain_Belgium/Etalle/wb/2018_complete_genome:0.0000005319):0.0000005319):0.0000224281,((KJ747406.1_UNVERIFIED__African_swine_fever_virus_isolate_Kashino_04/13_genomic_sequence:0.0001122679,(MK628478.1_African_swine_fever_virus_isolate_ASFV/LT14/1490_complete_genome:0.0000372327,MW306192.1_African_swine_fever_virus_isolate_ASFV/Ulyanovsk_19/WB-5699_complete_genome:0.0000858098):0.0000159564):0.0000057470,MT459800.1_African_swine_fever_virus_isolate_ASFV/Kabardino-Balkaria_19/WB-964_complete_genome:0.0000531907):0.0000005319);
